# Immunomodulatory Role of the Stem Cell Circadian Clock in Muscle Repair

**DOI:** 10.1101/2024.05.24.595728

**Authors:** Pei Zhu, Eric M Pfrender, Adam W T Steffeck, Colleen R Reczek, Yalu Zhou, Abhishek Vijay Thakkar, Neha R Gupta, Amber Willbanks, Richard L Lieber, Ishan Roy, Navdeep S Chandel, Clara B Peek

## Abstract

The circadian clock orchestrates vital physiological processes such as metabolism, immune function, and tissue regeneration, aligning them with the optimal time of day. This study identifies an intricate interplay between the circadian clock within muscle stem cells (SCs) and their capacity to modulate the immune microenvironment during muscle regeneration. We uncover that the SC clock provokes time of day-dependent induction of inflammatory response genes following injury, particularly those related to neutrophil activity and chemotaxis. These responses are driven by rhythms of cytosolic regeneration of the signaling metabolite NAD^+^. We demonstrate that genetically enhancing cytosolic NAD^+^ regeneration in SCs is sufficient to induce robust inflammatory responses that significantly influence muscle regeneration. Furthermore, using mononuclear single-cell sequencing of the regenerating muscle niche, we uncover a key role for the cytokine CCL2 in mediating SC-neutrophil crosstalk in a time of day-dependent manner. Our findings highlight a crucial intersection between SC metabolic shifts and immune responses within the muscle microenvironment, dictated by the circadian rhythms, and underscore the potential for targeting circadian and metabolic pathways to enhance tissue regeneration.

## Introduction

The circadian clock, the body’s intrinsic time-keeping mechanism, operates through a molecular clock in almost all cells within the body^1^. It comprises a central clock in the brain’s hypothalamic suprachiasmatic nucleus and peripheral clocks distributed throughout the rest of the body, influencing behaviors like sleep and physiological processes including core body temperature, hormone secretion, immune function, and metabolism^2,3^. Circadian rhythms ensure these activities align optimally with the time of day, maximizing fitness. On the molecular level, the clock functions as a transcription-translation feedback loop consisting of activators (CLOCK, BMAL1) which direct the expression of thousands of genes, including their own repressors (PER1-3, CRY1-2). Accessory proteins can stabilize the oscillation, including the nuclear receptors, REVERBa/b and RORa/b^1^. Previous work from our group^4^ and others^5^ uncovered that the efficiency of muscle repair is subject to time-of-day (TOD) variations, suggesting that circadian rhythms influence tissue regeneration. However, the mechanisms by which circadian clocks modulate this process have remained largely elusive.

Muscle regeneration is a highly coordinated, dynamic process involving intricate interactions between muscle stem cells (a.k.a. satellite cells, SCs) and the immune system^6^. SCs are crucial for muscle repair, activated by injury to proliferate and differentiate, thereby replenishing the muscle fiber pool^7^. Neutrophils, the first responders to injury, play a critical role in initiating repair by clearing debris and secreting cytokines and chemokines^8^, thus fostering an environment conducive to healing and influencing the behavior of SCs and other progenitor cells^9,10^. Nevertheless, the interactions between neutrophils and SCs, especially considering the metabolic changes triggered by muscle injury, are yet to be fully understood.

This study introduces a novel role for the SC circadian clock in modulating these interactions through metabolic responses to injury. We demonstrate that the SC clock drives the production of the oxidized form nicotinamide adenine dinucleotide (NAD^+^) via anaerobic glycolysis in a TOD-dependent manner. Using a genetic mouse model expressing the cytosolic NADH oxidase from Lactobacillus brevis (*LbNOX*) within SCs, we found that cytoplasmic NAD^+^ regeneration is sufficient to boost myofiber repair. Notably, we uncovered a previously unrecognized role for NAD^+^ regeneration in SCs in the activity of PARP1 and NFkB, extending beyond previously identified NAD^+^-dependent control of Sirtuins and myogenic gene deacetylation. We show that hypoxic NAD^+^ regeneration acts in SCs to drive neutrophil chemotaxis-related gene expression and enhance neutrophil infiltration and their interaction with SCs. In particular, the NAD^+^-regulated chemokine *Ccl2* exhibited varied expression levels across different times of day, influencing the quality and rate of muscle regeneration.

Our findings not only provide insights into the circadian regulation of muscle regeneration but also highlight a complex metabolic-immune interface within the muscle microenvironment, potentially explaining the compromised muscle regeneration observed in conditions characterized by disrupted circadian rhythms, such as obesity, diabetes, and aging. This study opens new avenues for therapeutic strategies targeting the interplay between circadian biology, metabolism, and immune responses to optimize tissue regeneration.

## Results

### NAD^+^-regenerating SCs express genes modulating neutrophil functions

Under conditions of hypoxia, pyruvate, the end product of glycolysis, is converted into lactate by lactate dehydrogenase A (Ldha). This process involves the reduction of pyruvate with electrons from NADH, facilitating the regeneration of NAD^+^ (Fig. 1A). Using the myotoxin cardiotoxin to injure mouse muscles, we observed that the injury creates an acute hypoxic environment within skeletal muscle, a condition confirmed by the substantial stabilization of HIF proteins in the initial days post-injury (Supplementary Fig. 1A). SCs within the muscle are also subjected to this hypoxic stress, as demonstrated by microarray analysis from a previous study^9^ that examined the temporal gene expression in SCs under both homeostatic conditions and after cardiotoxin-induced injury at various time points. Key HIF target genes, such as *Hk2*, *Vegf⍺*, and *Epo*, were markedly upregulated in SCs by day 2 post-injury, followed by a decline over time (Supplementary Fig. 1B). Notably, *Ldha* expression, known for its higher pyruvate affinity and preference for converting pyruvate to lactate^11^, was significantly increased during the hypoxia phase (Supplementary Fig. 1C). Conversely, its isoform *Ldhb*, which favors converting lactate back to pyruvate due to its higher affinity for lactate, was downregulated in the days immediately following the injury (Supplementary Fig. 1C). This suggests that SCs generate NAD^+^ through pyruvate fermentation in the early days post-myotoxin-induced muscle injury. We then performed RNA-sequencing on quiescent SCs (QSCs) from homeostatic muscles and the activated SCs (ASCs) from injured muscles on day 1 post-injury, a time when cytoplasmic NAD^+^ is predicted to be highly produced (Fig. 1B). The transcription profiles of SCs in these two states showed significant differences, with the first principal component accounting for more than 90% of the variance (Fig. 1C). Additionally, thousands of genes were differentially expressed (|LogFC| > 0.5, adj. p < 0.05) (Supplementary Fig. 2A). As anticipated, genes related to hypoxia and glycolysis signaling pathways were significantly upregulated in ASCs at 1-day post-injury (dpi) compared to QSCs (Fig. 1D), confirming the hypoxic state of these ASCs. Interestingly, genes associated with the inflammatory response, particularly those involved in neutrophil chemotaxis and migration, were also highly expressed in ASCs compared to QSCs (Fig. 1E). Given that neutrophils are the initial wave of immune cells to infiltrate the wounded site, peaking at day 1 post-injury^12^, these data suggest that ASCs play a role in modulating neutrophil activity at the early stages of muscle regeneration.

**Fig. 1.**
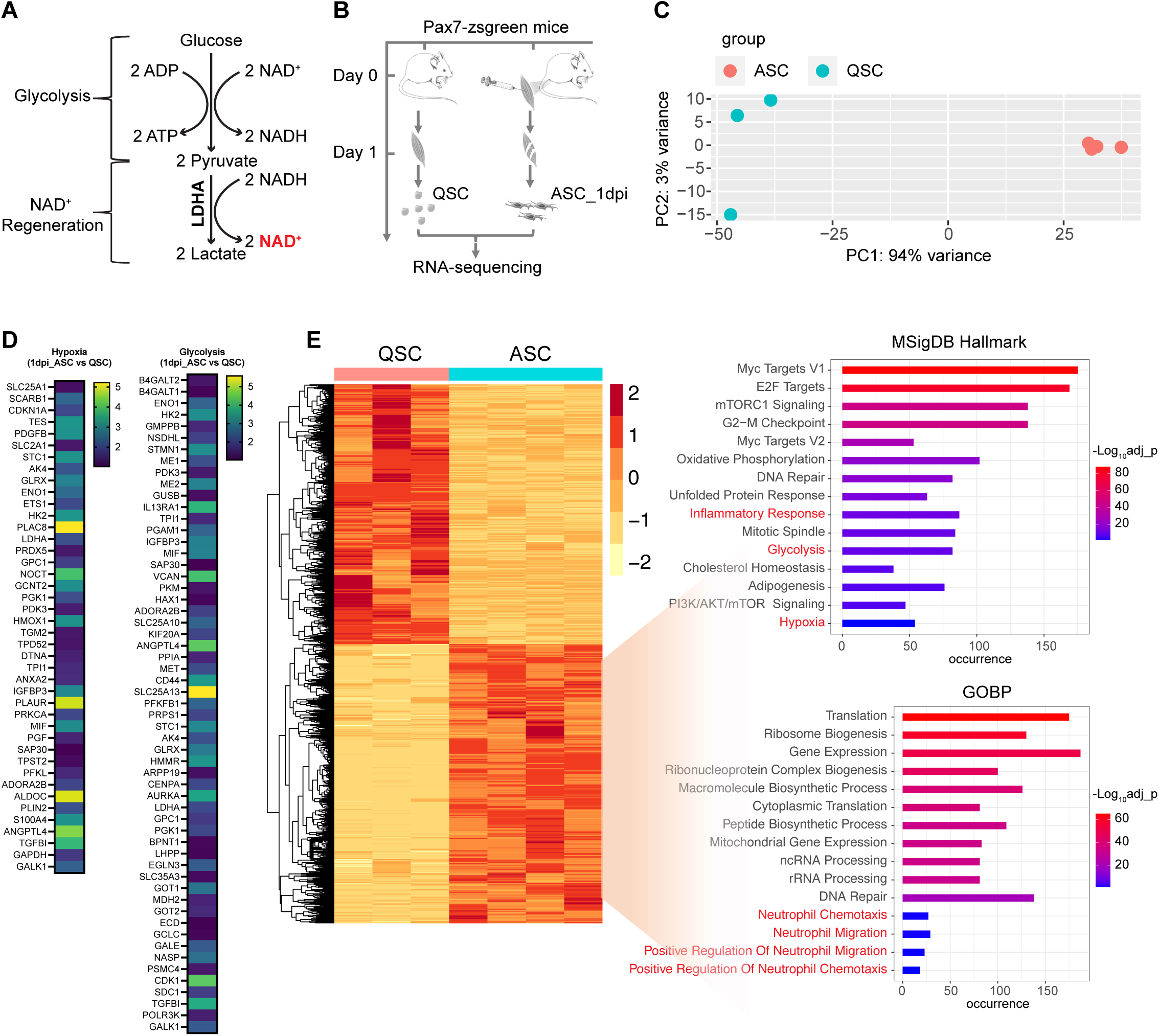
Elevated expression of glycolysis and inflammation-related genes in SCs under hypoxic conditions. (A) Schematic representation of the anaerobic glycolysis pathway. (B) Study design: SCs were extracted from the hindlimb muscles of Pax7-zsgreen mice, both uninjured and following cardiotoxin-induced injury, on day 1 post-injury, and subjected to RNA sequencing for transcriptome analysis. (C) Principal component analysis depicting the transcriptional disparities between ASCs at 1 dpi and QSCs. (D) Heatmap showing genes significantly upregulated (adjusted *p*-value < 0.05) in ASCs at 1 dpi, highlighting associations with hypoxia and glycolysis, in comparison to QSCs. The color scale represents the log2 fold change, indicating the magnitude of gene expression changes. (E) Pathway enrichment analysis, utilizing MSigDB hallmarks and gene ontology biological processes, for genes significantly upregulated (adjusted *p*-value < 0.05) in ASCs at 1 dpi relative to QSCs.

### SCs express genes facilitating neutrophil attachment, adhesion, and chemoattraction

Next, we performed single-cell RNA sequencing to explore the potential interactions between SCs and neutrophils during the early hypoxic stages of muscle regeneration using mononucleated cells collected from both intact and cardiotoxin-injured tibialis anterior (TA) muscles at days 1 and 3 post-injury (Fig. 2A). Following quality control procedures, the samples were integrated into a comprehensive transcriptomic atlas comprising 58,330 cells (Fig. 2B). We then performed unsupervised clustering and, through uniform manifold approximation and projection (UMAP), identified 21 cell types annotated based on the expression of established marker genes^13–15^ (Fig. 2B and Supplementary Fig. 3A). Notably, within this atlas, two distinct clusters expressing classical neutrophil markers (S100a8 and S100a9) were delineated, termed Neutrophils_1 and Neutrophils_2 (Fig. 2B and Supplementary Fig. 3A). Differential gene expression analysis revealed that Neutrophils_1 are more mature than the Neutrophils_2 subtype, as evidenced by a significantly higher expression of the chemokine receptor *Cxcr2*^16^ (Supplementary Fig. 3B). Correspondingly, gene ontology of biological process (GOBP) analysis showed that Neutrophils_1 exhibited elevated expression of genes associated with inflammatory responses (Supplementary Fig. 3C), signifying their pronounced inflammatory nature and advanced maturation. In contrast, Neutrophils_2 showed increased expression of genes related to lysosomal lumen acidification (Supplementary Fig. 3C), a characteristic of granulocyte differentiation of neutrophilic leukocyte precursors, which is essential for degradation of phagocytosed microorganisms at the inflammation sites^17^. Intriguingly, enrichment analysis using the literature-based network in the Elsevier Pathway Collection highlighted surface expression markers regulated by myoblasts in Neutrophils_1 and by myeloblasts in Neutrophils_2, respectively (Supplementary Fig. 3C). This observation underscores a potential modulatory role of SCs in neutrophil activity, particularly influencing Neutrophils_1 in the context of our experimental conditions.

**Fig. 2.**
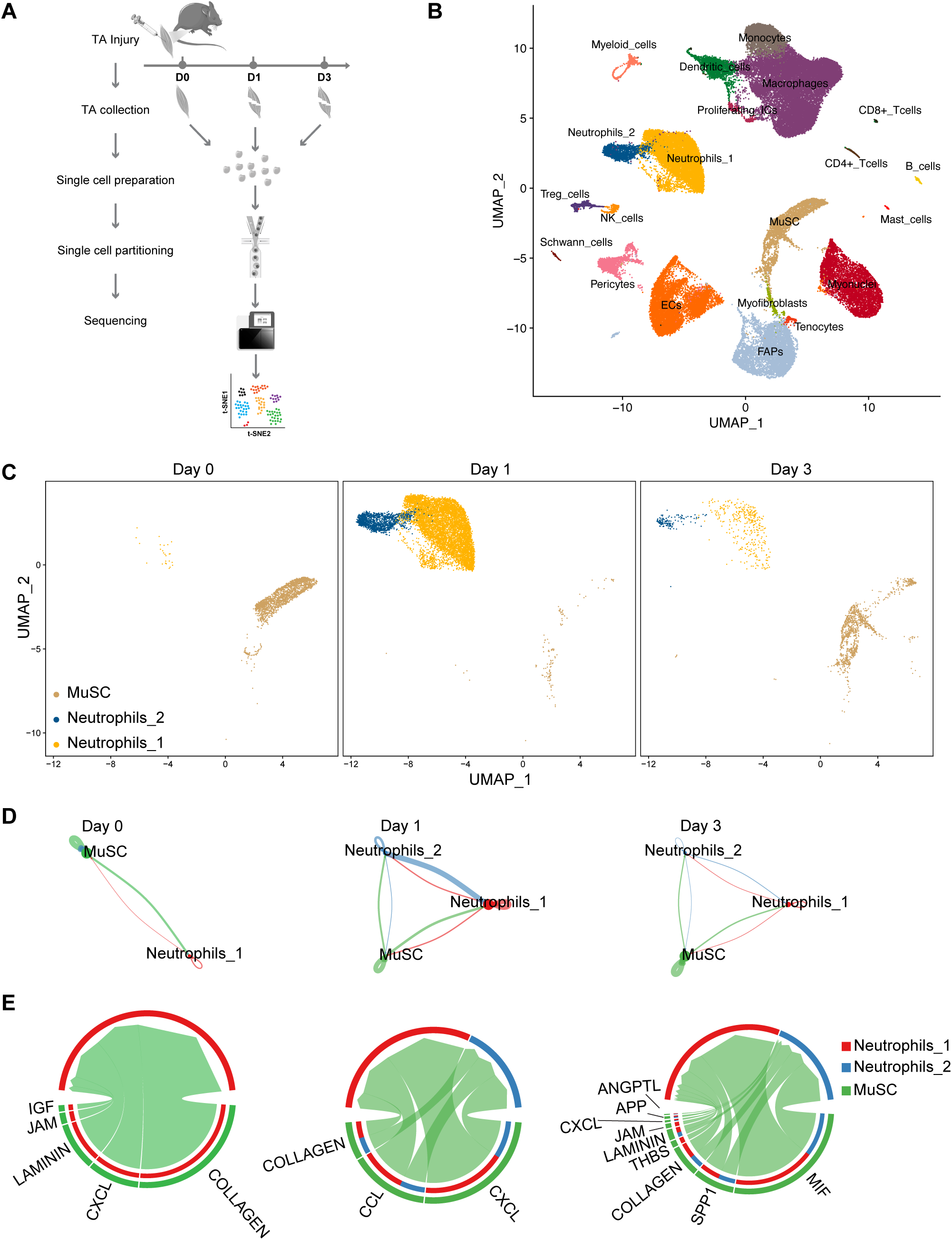
Interactions between SCs and neutrophils during early muscle regeneration. (A) Experimental design: mononucleated cells from the tibialis anterior (TA) muscles of both hindlimbs were collected at ZT4 and ZT16 on day 0, 1, and 3 after injury. Single cells were collected through fluorescence-activated cell sorting (FACS), excluding dead cells identified by propidium iodide (PI) staining. Samples from each time point and day were captured in droplet emulsions using a 10x Chromium Controller (10x Genomics), aiming for 10,000 cells per sample, and then sequenced. (B) UMAP visualization of 21 annotated cell types within pooled TA muscle cells from all 6 samples. EC, endothelial cells; FAPs, fibro-adipogenic progenitors. (C) UMAP of neutrophil and MuSC single-cell transcriptomes split by days post-injury. (D) Circle plot showing the aggregated cell-cell communication network among MuSCs and individual cluster of neutrophils at various days post-injury. (E) Chord diagram showing the specific ligand-receptor or signaling pathway-mediated interactions among the three cell clusters.

Based on our finding that neutrophil-related genes are induced at day 1 post-injury (Fig. 1E), we next focused on the regulatory interplay between MuSCs and neutrophils during early muscle regeneration. To this end, we spared neutrophils and MuSCs from the total cell populations and re-examined their dynamics over time using UMAP. A few Neutrophils_1 but no Neutrophils_2 were initially presented in homeostatic muscles (day 0) (Fig. 2C). There was a marked increase in both neutrophil types on day 1 post-injury, which then sharply declined by day 3 post-injury (Fig. 2C). Potential cross-communication between MuSCs and the two neutrophil types were delineated using CellChat, which can infer intercellular signaling network based on single-cell RNA sequencing data^18^. This was illustrated through vectorial circle plots representing interactions between any two cell groups (Fig. 2D). The communicative links mediated by MuSCs were most prominent on day 1 post-injury, highlighted by the densest edge weights directed from MuSCs toward the neutrophil populations (Fig. 2D). It is noteworthy that the nature of MuSC-initiated interactions evolved across the different days. Interactions involving laminin and collagen signaling, which facilitate cell attachment or adhesion, along with α chemokines featuring the CXC motif (CXCL) secreted by MuSCs, were prevalent at all observed time points. In contrast, β chemokines with the CC motif (CCL) were exclusively secreted by ASCs at 1 dpi, playing a crucial role in mediating MuSC communication with both neutrophil types (Fig. 2E).

### NAD^+^ augmentation induces chemokine gene expression and mediates neutrophil recruitment in muscle regeneration

CXCL and CCL chemokines are potent neutrophil chemoattractants^19^. Indeed, our bulk RNA sequencing (Fig. 1B) revealed that multiple chemokine genes of both families were significantly upregulated in ASCs at 1 dpi compared to QSCs (Fig. 3A), coinciding with increased NAD^+^ production during hypoxia. Given that hypoxia is known to modulate chemokine and chemokine receptor expression, thereby influencing tumor cell migration, invasion, and metastasis^20,21^, we investigated whether low oxygen tension or more specifically, anaerobically regenerated NAD^+^, could trigger chemokine gene expression in SCs. To uncouple NAD^+^ production from hypoxia, we overexpressed cytosolic NADH oxidase from *Lactobacillus brevis* (*LbNOX*) in myoblasts, which allows for a compartment-specific elevation of the NAD^+^/NADH ratio^22^. In our engineered *cytoLbNOX* mouse model, a CAG promoter-driven *cytoLbNOX* gene is inserted downstream of a loxP-flanked stop codon to preclude *LbNOX* expression prior to induction (Supplementary Fig. 4A). Cre recombinase-mediated excision of this stop codon activates *cytoLbNOX* and an internal ribosome entry site (IRES)-concatenated enhanced green fluorescence protein (EGFP) (Supplementary Fig. 4B). Primary myoblasts from *cytoLbNOX*-floxed mice were isolated and treated with Cre-expressing or control adenovirus (Fig. 3B). *CytoLbNOX*-overexpressing myoblasts exhibited a significant increase in intracellular NAD^+^ (Supplementary Fig. 4C), a decrease in NADH levels (Supplementary Fig. 4D), and a reduced NADH/NAD^+^ ratio (Supplementary Fig. 4E). Seventy-two hours post infection, cells were exposed to normoxic or hypoxic conditions for 24 hours. Echoing findings in cancer research, hypoxia significantly upregulated the candidate chemokines and integrins identified in ASCs at 1dpi (Fig. 3C). Remarkably, the presence of *cytoLbNOX*-sufficed to upregulate these genes, even in normoxia (Fig. 3C). Conversely, the application of Galloflavin, a potent lactate dehydrogenase inhibitor that impedes NAD^+^ regeneration (Supplementary Fig. 5A)^23^, significantly diminished the expression of these immunomodulator genes under hypoxia conditions (Supplementary Fig. 5B). Together, these data underscore the role of hypoxic NAD^+^ regeneration in the regulation of immune gene expression.

**Fig. 3.**
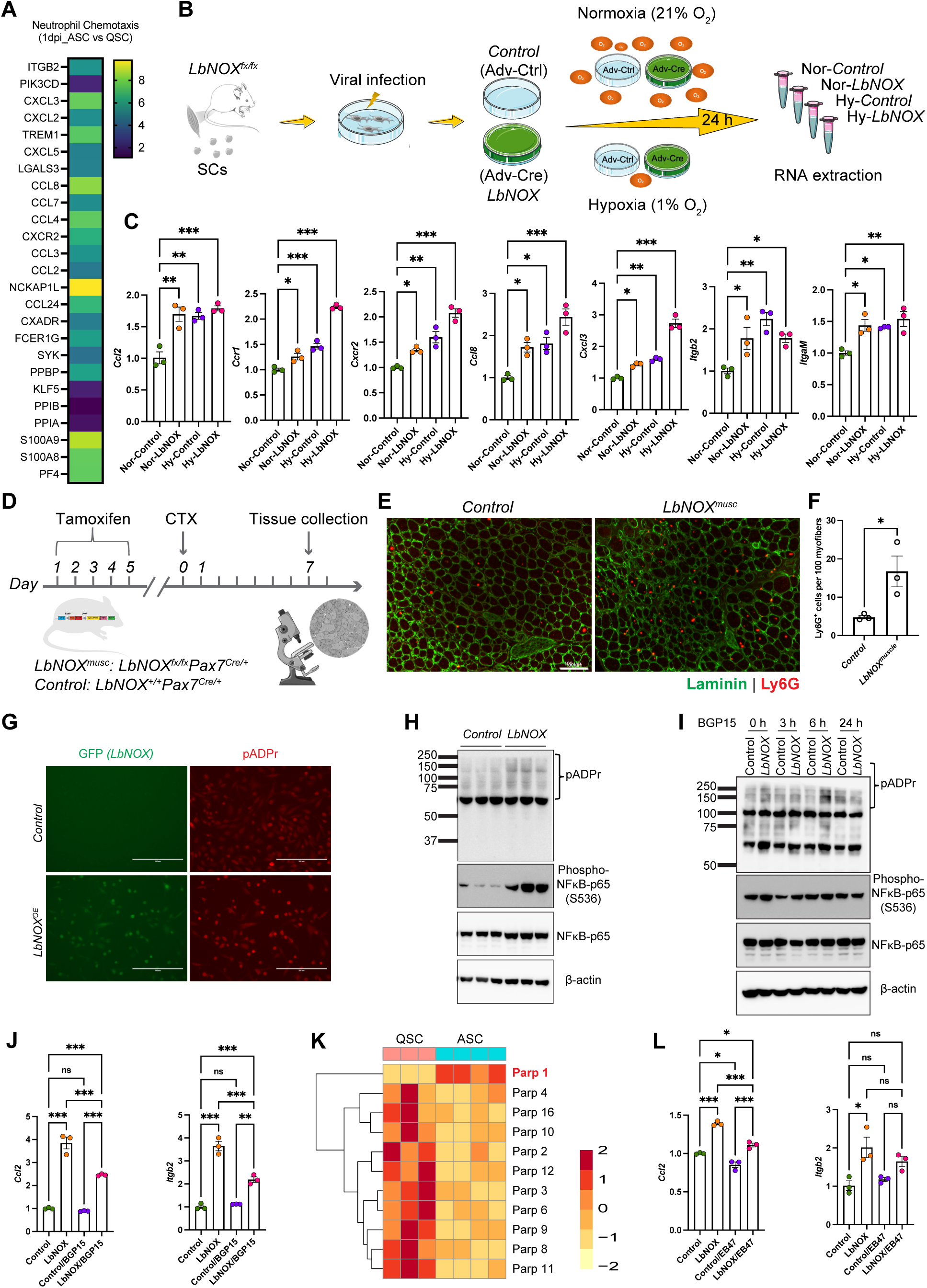
NAD^+^ augmentation induces immunomodulator expression via Parp-mediated PARylation and NK-κB activation. (A) Heatmap showing significantly upregulated immunomodulator genes (adjusted *p*-value < 0.05) in ASCs at 1 dpi compared to QSCs. Color scale denotes log2 fold change. (B) Experimental design: myoblasts from Cyto-*LbNOX^fx/fx^* mice were infected with an adenovirus expressing Cre recombinase (Adv-Cre) to induced *LbNOX* overexpression and cytosolic NAD^+^ augmentation. Adenovirus without Cre (Ade-Ctrl) served as the control. Three days after infection, both control and *cytoLbNOX*-overexpressing myoblasts were subjected to either hypoxia (1% O_2_) or normoxia (21% O_2_) for 24 hours, followed by RNA extraction and qPCR analysis for gene expression. (C) qPCR analysis of selected immunomodulator genes in control versus *LbNOX*-overexpressing myoblasts under conditions described in (B). * p < 0.05, ** p < 0.01, *** p < 0.001 by One-Way ANOVA, comparing each group to Ade-Ctrl-infected myoblasts under normoxia conditions. (D) Experimental design: MuSC conditional *LbNOX* expression (*LbNOX^musc^*) was achieved by crossing Cyto-*LbNOX^fx/fx^* mice with mice expressing Cre recombinase under the Pax7 promoter. Controls were heterozygous for the Pax7-cre allele without the *LbNOX* allele. *LbNOX* expression was induced via 5 consecutive intraperitoneal tamoxifen injections, with a 7-day recovery before cardiotoxin injury to TA muscles. Neutrophil retention was assessed 7 days post-injury by immunohistochemistry for Ly6G (n = 3 per group). (E) Representative immunohistochemistry images showing Ly6G^+^ neutrophils in TA muscle cryo-sections from Control and *LbNOX^musc^* mice, with Laminin outlining myofiber boundaries. Scale bar, 100 µm. (F) Quantification of neutrophils (Ly6G^+^) under conditions described in (E). * p < 0.05 by unpaired student’s t-test. (G) Representative immunofluorescence images showing global PARylation in myoblasts from Control and *LbNOX^musc^* mice, where Cre-induced *LbNOX* overexpression was marked by GFP co-expression. Scale bar, 200 µm. (H,I) Western blot analysis of global PARylation, total and serine 536 phosphorylated NF-κB p65 subunit, with β-actin as the internal control in *LbNOX-*overexpressing and Control myoblasts at 96 h post induction in the absence (H) or presence (I) of BGP15. (J) qPCR quantification of immunomodulator genes in Control and *LbNOX*-expressing myoblasts treated for 6 hours with or without BGP15. * p < 0.05, *** p < 0.001 by One-Way ANOVA analysis. Ns, non-significant. (K) Heatmap illustrating the expression profiles of Parp family members in ASCs at 1 dpi compared to QSCs. The color scale denotes Z-scores. (L) qPCR quantification of *Ccl2* expression in Control and *LbNOX*-expressing myoblasts following a 6-hour treatment with or without EB47. * p < 0.05, *** p < 0.001 by One-Way ANOVA.

Next, using myoblasts exposed to both *cytoLbNOX* and hypoxia, we further assessed the contribution of NAD^+^ regeneration to hypoxia-induced immunomodulator gene expression. Two-way ANOVA analysis revealed that among the immunomodulator genes induced with both *cytoLbNOX* overexpression and hypoxia, the interaction effect between oxygen concentration and *cytoLbNOX*-mediated NAD^+^ regeneration was significant for genes including *Ccl2* (p = 0.008), *Itgb2* (p = 0.033), *Cxcl3* (p = 0.047), and *Ccr1* (p = 0.026). Even though significant, Cxcl3 and Ccr1 were more robustly induced in *cytoLbNOX*-expressing myoblasts exposed to hypoxia compared to those not expressing *cytoLbNOX* (Fig. 3C). This pattern was similar to other tested genes, such as *Ccl8* (p = 0.869), *Cxcr2* (p = 0.514), and *ItgaM* (p = 0.208), which showed a non-significant interacting effect between oxygen tension and *cytoLbNOX*-mediated NAD^+^ regeneration according to two-way ANOVA. This suggests that additional mechanisms might underlie their induction under hypoxic conditions beyond NAD^+^ regeneration. Importantly, for *Ccl2* and *Itgb2*, hypoxia treatment did not further induce increase of gene expression in myoblasts overexpressing *cytoLbNOX* (Fig. 3C), suggesting that hypoxia induces expression via NAD^+^ regeneration. Consistent with this observation, the addition of Galloflavin failed to impede their upregulation in *cytoLbNOX*-overexpressing myoblasts, further supporting the pivotal role of NAD^+^ regeneration, as opposed to lactate, in this context (Supplementary Fig. 5C).

Considering the chemoattractant properties of chemokines, we examined whether *cytoLbNOX*-driven NAD^+^ augmentation could enhance neutrophil recruitment within regenerating muscles *in vivo*. To this end, we crossed *cytoLbNOX-*floxed mice with a line expressing Cre recombinase under the *Pax7* promoter, resulting in muscle stem cell-specific *cytoLbNOX* expression (*LbNOX^musc^*). Following tamoxifen induction and cardiotoxin-induced TA muscle injury, we assessed neutrophil presence at 7 dpi via Ly6G immunohistochemistry (Fig. 3D). As expected, neutrophil numbers in control mouse TA muscles had largely diminished by 7 dpi (Fig. 3E). In contrast, *LbNOX^musc^* mice retained significantly more neutrophils than controls (Fig. 3E and 3F), aligning with our hypothesis that cytoplasmic NAD^+^ mediates neutrophil activity in regenerating muscle.

NAD^+^ serves as a critical co-substrate for two main families of post-translational modification enzymes: the adenosine diphosphate (ADP)-ribosyltransferase (ARTs) and the sirtuins (SIRTs)^24–26^. Interestingly, poly ADP-ribose polymerase (PARP) enzymes have been shown to activate the major immunoregulatory transcription factor NF-κB in macrophages (PMID: 10449410). Therefore, we asked whether cytoplasmic NAD^+^ can mediate neutrophil-specific chemokines via control of PARP and PARylation. Immunofluorescence analysis revealed a global elevation in PARylation in *cytoLbNOX*-overexpressing myoblasts relative to controls (Fig. 3G). This was substantiated by Western blot analysis, which detected a significant increase in total cellular poly-ADP-ribosylated proteins in these myoblasts (Fig. 3H). This rise in PARylation coincided with NF-κB activation, as evidenced by the enhanced phosphorylation of the NF-κB p65 subunit at serine 536 (Fig. 3H). The increase in protein PARylation and NF-κB activation induced by NAD^+^ overproduction mirrored those observed in myoblasts subjected to hypoxia (Supplementary Fig. 5D). Notably, treatment with BGP15, a broad PARP inhibitor^27^, not only reduced intracellular protein PARylation (Fig. 3I) but also effectively diminished NF-κB activation, as evidenced by the significantly decreased phosphorylation of the NF-κB p65 subunit at serine 536 in *cytoLbNOX*-overexpressing myoblasts (Fig. 3I). As a result, the expression of *Ccl2* and *Itgb2* was notably decreased by this inhibitor in myoblasts expressing *cytoLbNOX*, whereas no significant effect was observed in control cells (Fig. 3J). This implies that PARP-mediated PARylation contributes to the upregulation of neutrophil chemotaxis-related gene expression in myoblasts with elevated NAD^+^ levels through the activation of the NF-κB signaling pathway. To identify the specific *Parp* gene involved in this NAD^+^-induced PARylation, we compared the expression of *Parp* family genes in ASCs at 1dpi versus QSCs, and found that *Parp1* was the only family member significantly upregulated in 1dpi ASCs compared to QSCs (Fig. 3K), a finding corroborated by a peak at 2 dpi and a return to baseline over time in previously published transcriptomic data from MuSCs isolated at various time points following cardiotoxin injury (Supplementary Fig. 2B)^9^. Finally, EB47, a PARP1-specific inhibitor ^28^, significantly suppressed gene expression related to neutrophil chemotaxis in *cytoLbNOX*-overexpressing myoblasts, akin to the effects of BGP15, but not in control cells (Fig. 3L). Taken together, these data indicate the involvement of PARP1 in the NAD^+^-dependent regulation of neutrophil signaling from activated SCs.

### Hypoxic regulation of neutrophil immunomodulatory gene expression is under circadian control

Given the established role of the SC circadian clock in regulating hypoxic NAD^+^ production in myoblasts and muscle repair capacity following injury^4^, we hypothesized that NAD^+^-induced immunomodulatory gene expression might also be under circadian control. In support of this, total RNA sequencing of wildtype and *Bmal1^-/-^* primary myoblasts exposed to acute hypoxia (6 hours, 1% O_2_) (Supplementary Fig. 6A) revealed a marked downregulation of chemokine signaling pathway-related genes (Supplementary Fig. 6B and S6C). Next, we performed cardiotoxin-induced injuries in TA muscles at Zeitgeber (ZT) 4 and ZT16, corresponding to periods of low and high BMAL1 activity, respectively^4,29^. ASCs were isolated at 24 hours (1 day)-post injury (Fig. 4A) and subjected to total RNA sequencing (Fig. 4B). Aligning with our previous findings that the HIF signaling pathway is modulated by circadian clock^4,30^, genes related to the hypoxia response were significantly more upregulated in ASCs from muscles injured at ZT16 than at ZT4 (Fig. 4C). Notably, amongst the genes specifically induced in ASCs following injury at ZT16 versus ZT4 were those associated with inflammatory response, particularly neutrophil chemotaxis and chemokine-mediated signaling pathways (Fig. 4C).

**Fig. 4.**
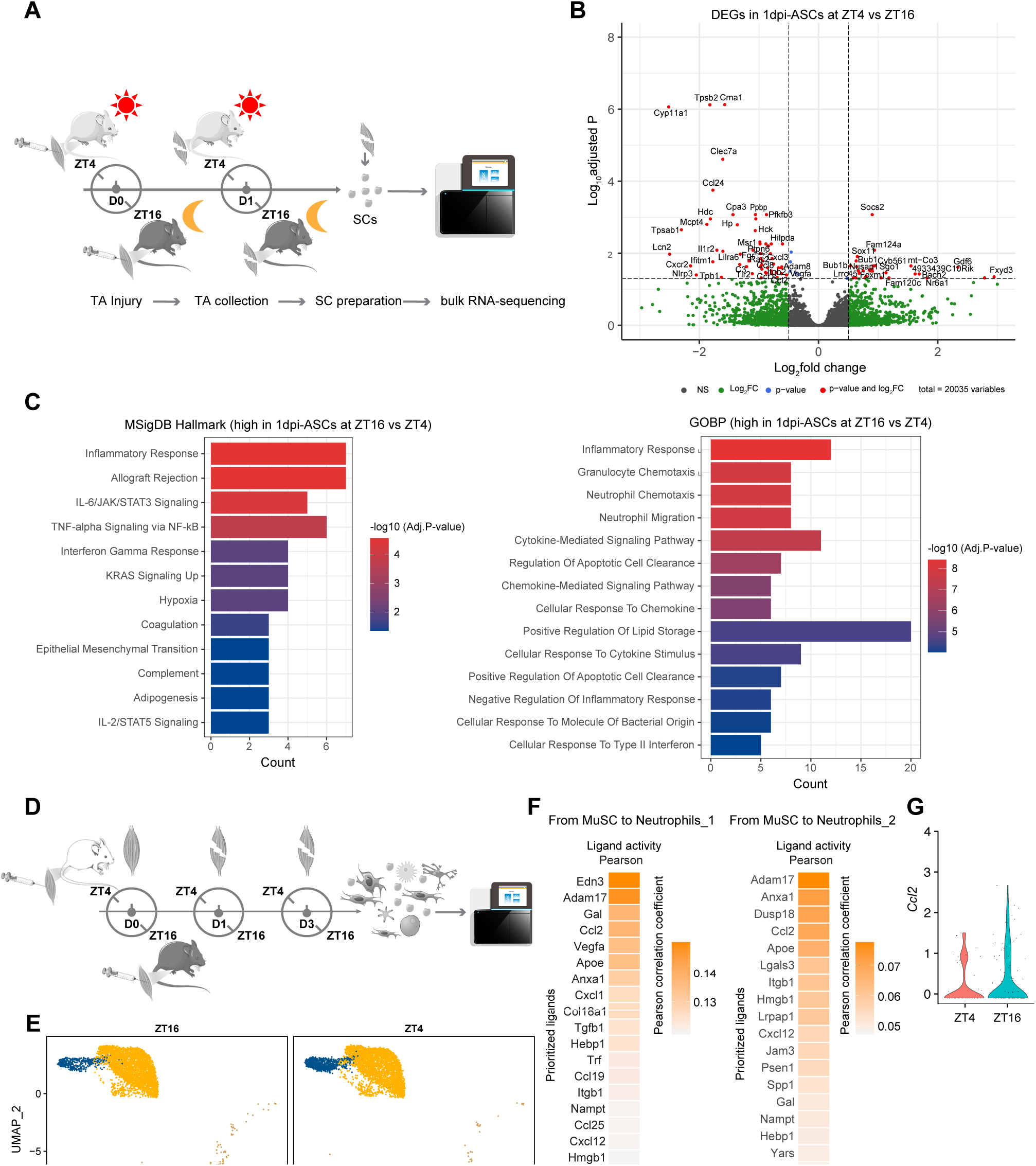
TOD-dependent immunomodulator expression in ASCs at day 1 post-injury. (A) Experimental design: Pax7-zsgreen mice were subjected to cardiotoxin-induced injury in the TA muscles at ZT4 and ZT16. Zsgreen+ SCs were isolated by FACS at the corresponding time points on day 1 post-injury and analyzed using RNA sequencing. (B) Volcano plot showing differentially expressed genes (adjusted p < 0.05) in ASCs harvested from injured TA muscles at ZT4 compared to ZT16 at day 1 post-injury. (C) Signaling pathway enrichment analysis for genes more highly expressed in ASCs at ZT16 compared to ZT4 at 1 dpi. (D) Experimental recap: this panel reiterates the experimental setup detailed in Fig. 2A, focusing on the timing of injury and subsequent collection of mononucleated cells at ZT4 and ZT16. (E) UMAP of neutrophil and MuSC single-cell transcriptomes split by the time of day on day 1 post-injury. (F) Heatmap of ligand activities by the NicheNet analysis showing ranked potential ligands expressed in the signal ‘sender’, MuSC, that could predict the observed differentially expressed genes in the indicated neutrophil populations between ZT4 and ZT16 on day 1 post-injury. (G) Violin plot comparing expression levels of *Ccl2* in ASCs at ZT4 and ZT16 on day 1 post-injury.

Our mononucleated muscle cell samples for single-cell RNA sequencing described earlier were collected at both ZT4 and ZT16 from both homeostatic TA muscles and those injured at ZT4 and ZT16 on 1 and 3 dpi (Fig. 4D). Re-examining the time-of-day (TOD)-dependent dynamics within the UMAP encompassing neutrophils and MuSCs (Fig. 4E), we employed NicheNet^31^ to dissect TOD-dependent communication differences between MuSCs and neutrophils, designating MuSCs as the signal ‘sender’ and neutrophils as the ‘receiver’. According to NicheNet’s guidelines, a Pearson correlation coefficient score around 0.10 or higher indicates significant ligand-mediated crosstalk. Our results showed more substantial ligand-mediated interactions between MuSCs and the Neutrophils_1 population, while interactions with Neutrophils_2 did not achieve a high Pearson score (Fig. 4F). Interestingly, *Ccl2* emerged as the sole CCL-type chemokine among the top ligands, exhibiting high activity in inducing target gene expression in Neutrophils_1 (Fig. 4F and S7). Furthermore, *Ccl2* gene expression was notably higher in ASCs at ZT16 compared to ZT4 on 1 dpi (Fig. 4G and 3A).

### *Ccl2* expression in SCs is associated with the TOD-dependent differences in SC proliferation

A previous study demonstrated that conditioned medium from neutrophil cultures can promote myoblast proliferation while suppressing their differentiation^32^. This prompted us to investigate whether the TOD-dependent interactions between MuSCs and neutrophils might reciprocally influence SC fate decisions during muscle regeneration. To explore this, we analyzed the cellular heterogeneity within the MuSC populations from our single cell-transcriptomic atlas. Unsupervised clustering identified 7 distinct subpopulations of myogenic cells (Fig. 5A). Notably, cells in cluster 3 exhibited high expression of *Pax7* and genes like *Spry1*, *Cd34*, *Cdkn1b*, and *Pdk4*, which are recognized markers of deep quiescent SCs (dQSCs)^33–35^. Further classification of the clusters was based on gene sets indicative of early commitment (e.g., *Myf5*, *Egr2*, *Egr3*, *Myc*, *Jund*, *Dnajb1*, *Ccl8*)^34,35^, activation/proliferation (*Myod1*, *Mki67*, *Cdk1*, *Ccnb1*)^36^ and differentiation (*Myog*, *Myl4*)^7,37^, leading to their designation as primed QSC 1 and 2, committed SCs, dividing SCs, differentiating SCs, and renewing SCs (Fig. 5A and Supplementary Fig. 8A). A temporal analysis on UMAP highlighted that quiescent SCs and some committed SCs were prevalent in homeostatic muscles, whereas dividing and differentiating SCs were mainly observed in regenerating muscles by 3 dpi (Supplementary Fig. 8B).

**Fig. 5.**
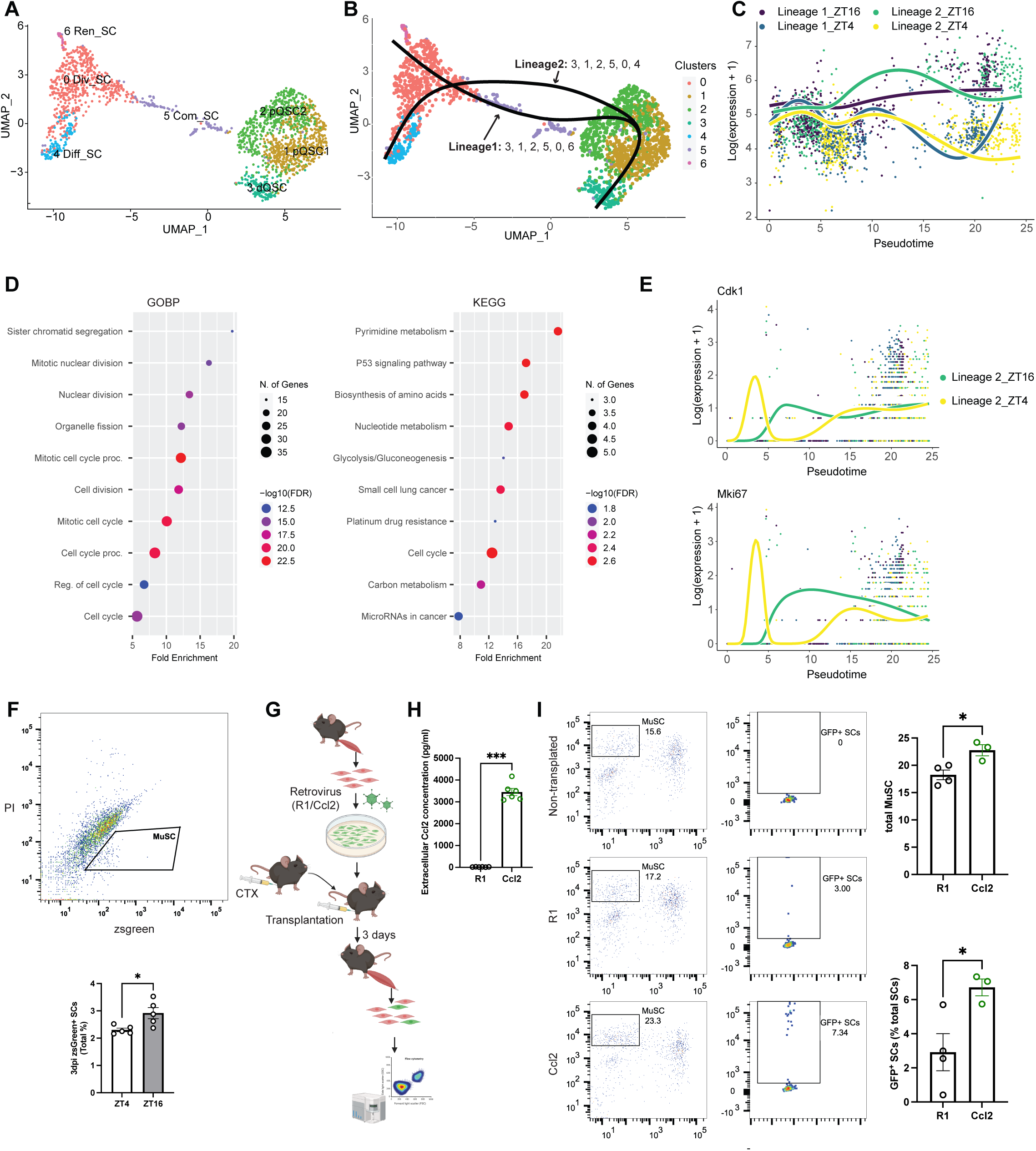
Ccl2 Secretion by SCs promotes TOD-dependent proliferative differences. (A) UMAP visualization of the subclustered MuSC populations, with subclusters defined as deep quiescent SCs (dQSC), primed quiescent SCs (pQSC). committed SCs (Com_SC), dividing SCs (Div_SC), differentiating SCs (Diff_SC), and renewing SCs (Ren_SC). (B) Pseudotime trajectory inference analysis of MuSC subclusters by Slingshot, with dQSCs serving as the starting point of pseudotime. (C) Pseudotime ordered single-cell expression trajectories for the genes that were most significantly differentially expressed between the time points of ZT4 and ZT16 in lineage 1 and 2. (D) Signaling pathway enrichment analyses on the differentially expressed genes from (C). (E) Pseudotime ordered single-cell expression trajectories for the cell cycle regulators Cdk1 and Mki67 at ZT4 and ZT16 within lineage 2. (F) Representative FACS plot for MuSC gating (zsGreen^+^) at 3 dpi (top panel) and quantification of MuSC frequency within total mononucleated muscle cells (bottom panel). * p < 0.05 by unpaired student’s t test (n = 5 mice per time point). (G) Experimental design: SCs from C57BL/6 wildtype mice were transduced with a retrovirus expressing GFP and either Ccl2 (Ccl2-virus) or a control virus (R1). Recipient mice (n = 4) then received bilateral TA muscle injuries via cardiotoxin injection. At 6 hours post-injury, 10^5^ transduced SCs were transplanted into each TA muscles, with Ccl2-virus infected SCs in one side and R1-virus infected SCs in the other. On day 3 post-transplantation, the total MuSC frequency and the percentage of GFP^+^ SCs within the total MuSC population in the recipient TA muscles were assessed by flow cytometry. (H) ELISA measurement of Ccl2 secretion in the media after 24 hours of culture from an equal number of myoblasts infected with either Ccl2 or control retrovirus. (I) Quantification of total MuSCs and GFP+ SCs in the recipient TA muscles as described in (G). * p < 0.05 by unpaired student’s t test.

To better understand the hierarchical organization and validate the classification of these myogenic subclusters, we applied Slingshot^38^, a single-cell trajectory inference analysis, to the entire myogenic cell population within the atlas. This analysis delineated a sequential trajectory from deep quiescence through primed quiescence and commitment/activation, leading to division, which then bifurcated toward self-renewal (lineage 1) or differentiation (lineage 2) (Fig. 5B). Along this pseudotime trajectory, *Pax7* expression gradually decreased, while *Myod1* expression increased in both lineages (Supplementary Fig. 8C). In the self-renewal lineage, *Myog* expression spiked before sharply decreasing, whereas in the differentiation lineage, *Myog* expression steadily rose (Supplementary Fig. 8C).

To elucidate differential gene expression patterns between ZT4 and ZT16 within each myogenic lineage, we employed the ‘conditionTest’ function from the tradeSeq package^39^. This analysis revealed 88 genes with significant differential expression (adjusted p-value < 0.05, fold change > 1) across pseudotime (Fig. 5C). Subsequent KEGG pathway enrichment analysis linked these genes predominantly to the cell cycle (Fig. 5D). We further assessed the expression trajectories of two key cell division markers, Cyclin-dependent kinase 1 (*Cdk1*) and *Mki67*, over pseudotime at both ZT4 and ZT16. Notably, within the differentiation lineage (lineage 2), ZT4 myogenic cells displayed an early peak in the expression of both regulators at lower pseudotime values-a peak not observed in ZT16 myogenic cells (Fig. 5E). Following this initial surge, expression levels in ZT4 cells plummeted to a baseline before experiencing a modest resurgence at higher pseudotime values. Conversely, in ZT16 cells within the same lineage, expression levels of *Cdk1* and *Mki67* ascended rapidly and remained consistently elevated throughout the observed pseudotime span (Fig. 5E).

The expression dynamics we observed suggest a potential regulatory effect of the TOD on key cell cycle regulators, which may in turn influence the proliferation trajectory of SCs after activation. To investigate this influence, we induced injury in TA muscles at ZT4 and ZT16 and then quantified the frequency of SCs in the total muscle cell population by day 3 post-injury, a peak time for dividing myonuclei^40^, using *Pax7*-zsgreen reporter mice^41^. Flow cytometry analysis revealed a significantly higher frequency of zsgreen^+^ SCs in TA muscles injured at ZT16 compared to those injured at ZT4 at 3 dpi (Fig. 5F).

To determine the extent to which SC-expressed CCL2 contributes to this TOD-dependent difference in SC proliferation, we transduced freshly isolated SCs with a *Ccl2*-expressing retrovirus and transplanted them into TA muscles of wildtype C57BL/6 mice 6 hours following cardiotoxin injury (Fig. 5G). The contralateral TA muscles were subjected to the same pre-injury but received control virus (R1)-transduced SCs. The transplanted myogenic progenitors, labeled by GFP carried by the viral vector, allowed for the tracking of GFP^+^-SCs within the total SCs population, characterized as VcamI^+^Scal1^-^CD31^-^CD45^-42^, at 3 dpi (Fig. 5G). Prior to this experiment, we first verified the role of CCL2 as a secreted chemokine by comparing its levels in culture media from myoblasts infected with either *Ccl2*-expressing or control retroviruses using ELISA (Fig. 5H). Notably, we observed a significantly higher frequency of *Ccl2*-overexpressing SCs in the TA muscles of recipient mice compared to those transplanted with control virus-infected SCs (Fig. 5I). This increase likely resulted from the CCL2 released into the niche by the exogenous SCs, which was also reflected in a significant increase in the total MuSC population compared to muscles receiving control virus-infected SCs (Fig. 5I).

### Neutrophil depletion abrogates TOD effects on SC dynamics and muscle regeneration

Our previous studies, along with others, showed a TOD-dependent variation in skeletal muscle repair following cardiotoxin-induced injury, with more effective regeneration occurring during active phase compared to the rest phase^4,5^. The enduring impact of this TOD effect remains underexplored. To investigate whether the difference is transient or has long-lasting consequences, we induced TA muscle injuries with cardiotoxin at ZT4 or ZT16, repeating the process three times at 30-day intervals (Fig. 6A). Histological examination conducted 30 days after the last injury revealed that TA muscles subjected to repeated injuries at ZT4, corresponding to a period of lower BMAL1 activity, exhibited a significantly higher proportion of small regenerated myofibers and lower proportion of large myofibers compared to those injured at ZT16 (Fig. 6B and 6C). Additionally, we assessed muscle strength by measuring torque under continuous electronic stimuli at multiple frequencies (Fig. 6D). Max tetanic force production in the uninjured TA muscle exhibited variations depending on the time of day, with increased peak force during tetanic contractions (Max tetanic) observed in the early rest phase (ZT4) compared to the active phase (ZT16) (Fig. 6E). We also assessed recovery of muscles following repeated injuries at either ZT4 or ZT16 by comparing the Max tetanic of injured muscles to that of uninjured ones at identical times of day. While the interaction between TOD and injury status did not reach significance by two-way ANOVA, we observed a trend towards reduced recovery in mice injured at ZT4, as compared to ZT16 (Fig. 6E). These findings point towards a potential lasting influence of TOD on muscle health and regeneration.

**Fig. 6.**
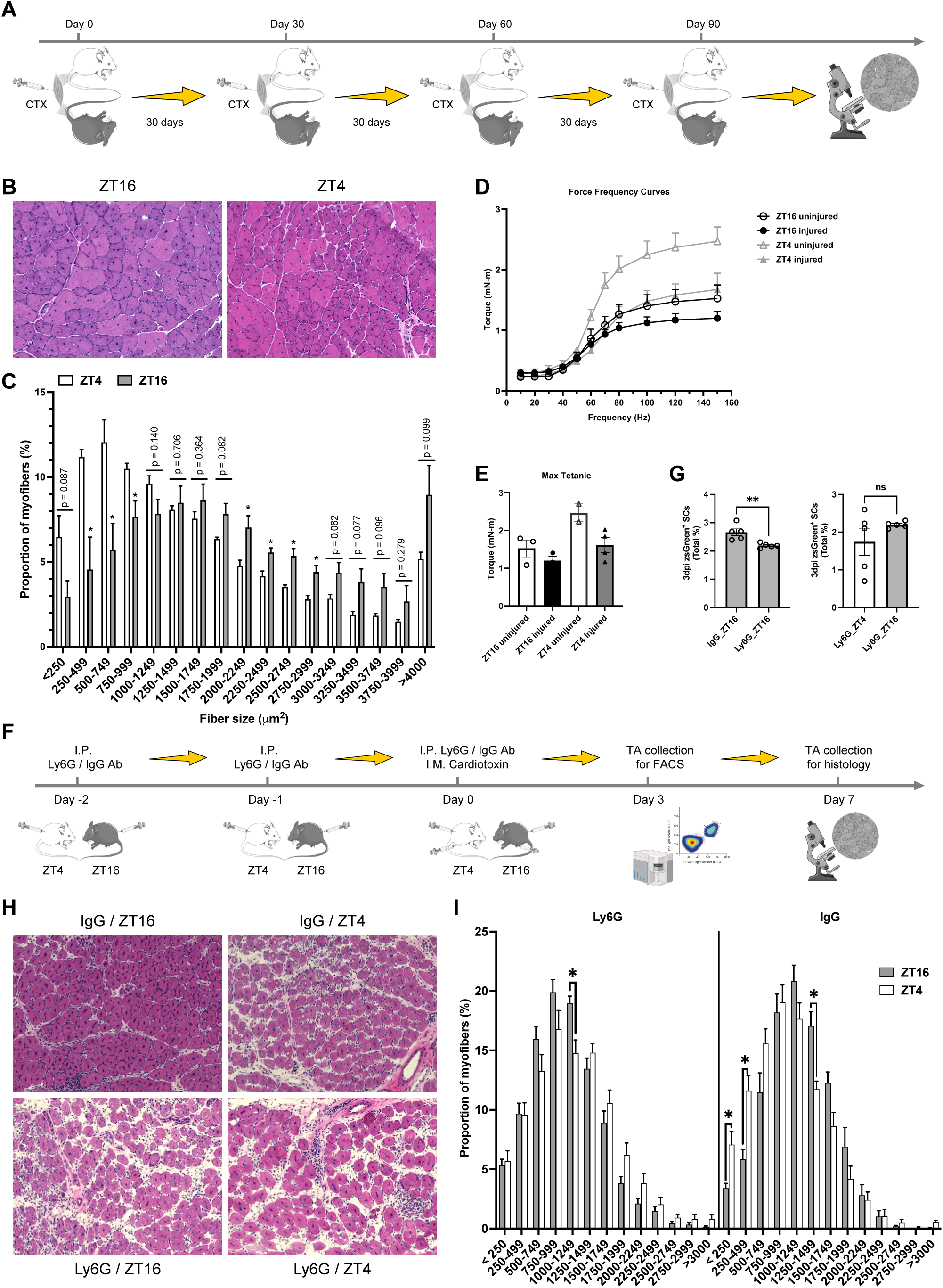
Reduction of TOD-dependent variations in SC proliferation and muscle regeneration through neutrophil depletion. (A) Experimental design: C57BL/6 wildtype mice (n = 3 per time point) were subjected to repeated cardiotoxin-induced injuries to their TA muscles at ZT4 and ZT16, thrice with a 30-day recuperation period between injuries. Thirty days post-final injury, TA muscles were extracted at their respective injury timings for muscle remodeling assessment by histology. (B) Representative H&E staining of the repeatedly injured TA muscles at ZT16 and ZT4. (C) Quantification of myofiber size distribution in the injured TA muscles depicted in (B). (D) Measurement of torque variations in response to different stimulation frequencies in the TA muscles that were repeatedly injured at ZT4 and ZT16. The torque at each frequency along the force frequency curve plotted as mean +/- SEM. (E) Peak torque generated in the force frequency curve (mean; SEM) for each group: ZT16u (1.526; 0.2237), ZT16i (1.200; 0.1101), ZT4u (2.470; 0.2365), ZT4i (1.614; 0.1968). (F) Experimental design: Pax7-zsGreen mice (n = 5 per group) underwent transient neutrophil depletion through 3 consecutive intraperitoneal injections of Ly6G antibody. Control mice received IgG antibody injections from the same host species as the Ly6G antibody. Following the final antibody injection, mice received intramuscular cardiotoxin injections in the TA muscles at ZT4 and ZT16. SC proliferation was then assessed by flow cytometry on 3 dpi, and muscle regeneration was evaluated via histology on 7 dpi at the corresponding injury timings. (G) Quantification of zsGreen^+^ SCs on 3 dpi in the injured TA muscles from both IgG and Ly6G-treated mice (top panel), and in TA muscles from neutrophil-depleted mice injured at ZT4 and ZT16 (bottom panel). ** p < 0.01 by unpaired student’s t-test. Ns, non-significant. (H) Representative H&E staining of TA muscles from IgG and Ly6G treated mice that were injured at ZT4 and ZT16. (I) Quantification of myofiber size distribution in the injured TA muscles shown in (H). * p < 0.05 by multiple *t* test.

To further probe the role of neutrophils in mediating the TOD effect on muscle regeneration, we depleted neutrophils in the early stages post-injury. Depletion was achieved through three successive intraperitoneal injections of a Ly6G antibody. On the day of the final injection, we induced TA muscle injuries with cardiotoxin and examined neutrophil infiltration 18 hours later (Supplementary Fig. 9A). Flow cytometry analysis confirmed that the Ly6G antibody specifically reduced neutrophil infiltration compared to the control IgG antibody, with minimal impact on other immune cells like monocytes (Supplementary Fig. 9B). Leveraging this neutrophil depletion strategy, we evaluated its effect on SC proliferation at 3 dpi and on muscle regeneration at 7 dpi, following cardiotoxin-induced injuries at ZT4 and ZT16 (Fig. 6F). Flow cytometry-based cell quantification showed that neutrophil depletion significantly reduced the frequency of SCs in TA muscles injured at ZT16 compared to those treated with control IgG antibody (Fig. 6G top panel). Importantly, neutrophil depletion eliminated the TOD-dependent disparities in SC proliferation at 3 dpi (Fig. 6G bottom panel) and in muscle regeneration at 7 dpi (Fig. 6G and 6I), suggesting that neutrophils are key modulators of the TOD effect in muscle repair processes.

### Concurrent activation of glycolysis and inflammatory pathways across muscle cell populations under hypoxia conditions

Our comprehensive analysis under hypoxic conditions unveiled a concurrent enhancement of anaerobic glycolysis and inflammatory responses in SCs, suggesting a potential systemic response mechanism to hypoxia. Utilizing GeneMANIA^43^, we established a functional link between the key glycolytic gene *Ldha* and neutrophil chemotaxis-associated genes upregulated in ASCs at 1dpi (Fig. 7A), suggesting this dual activation may be a broader cellular response to hypoxia.

**Fig. 7.**
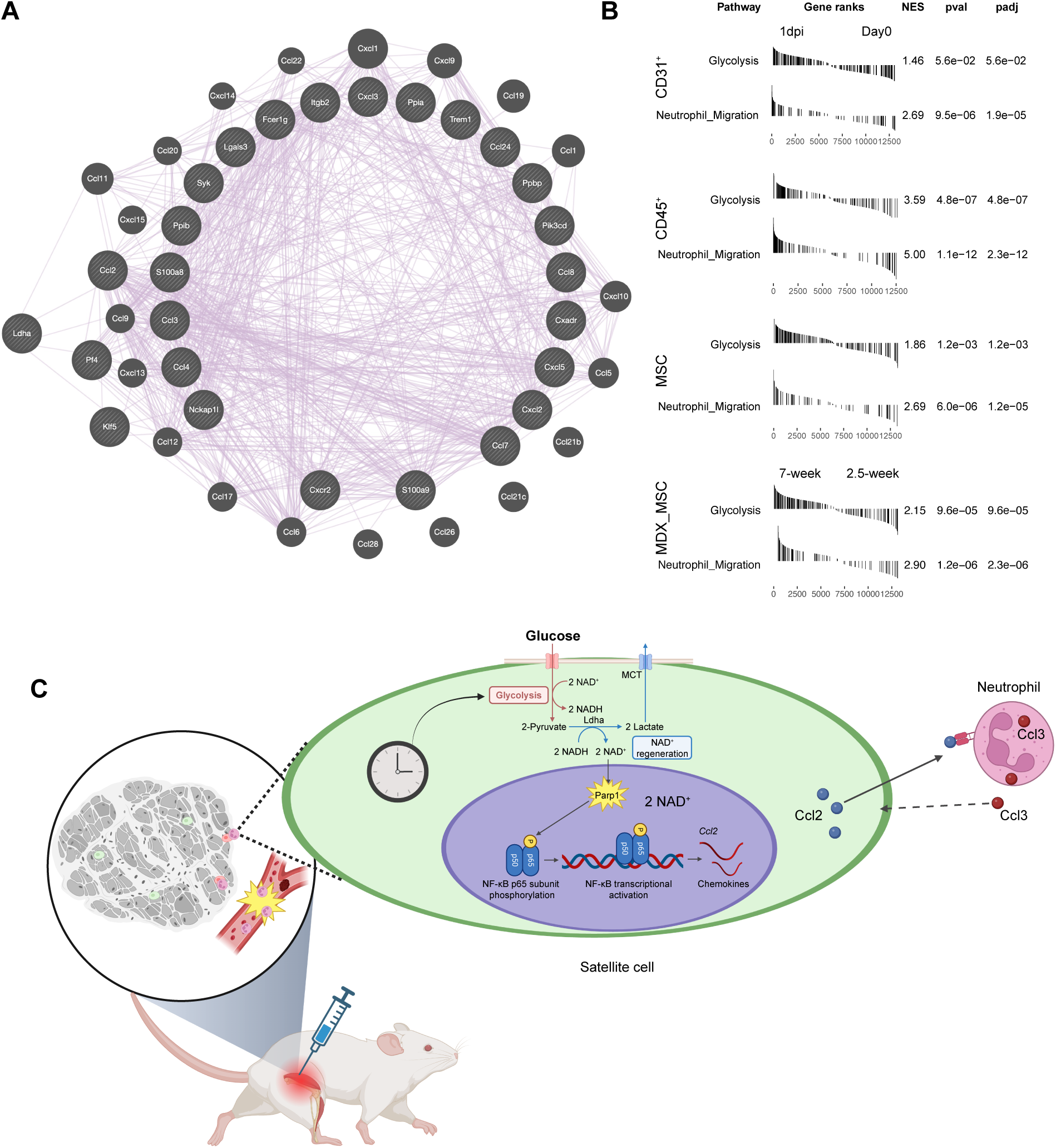
Co-activation of glycolysis and inflammatory signaling in muscle resident cells under hypoxic conditions. (A) a GeneMANIA gene interaction network showing gene-gene interactions based on co-expression among LDHA and genes associated with neutrophil chemotaxis and migration. (B) Fast preranked gene set enrichment analysis (fgsea) showing a positive correlation between pathways associated with glycolysis and neutrophil migration in resident cells within cardiotoxin-injured muscles at 1 dpi, compared to uninjured muscles, as well as in mesenchymal stromal cells (MSC) in muscles from 12-week-old versus 2.5-week-old *mdx* mice. (C) Mechanistic schematic depicting the process underlying TOD-dependent muscle regeneration. Following injury, skeletal muscle experiences passive hypoxia, triggering enhanced anaerobic glycolysis in SCs. The regeneration of cytoplasmic NAD^+^ during this process acts as a substrate as activated PARP1, which then activates NF-κB through the phosphorylation of the p65 subunit at serine 536. This activation further promotes the expression of immunomodulator genes (e.g., *Ccl2*), aligning with a peak in neutrophil infiltration into the wounded site. This synchronization facilitates interactions between SCs and neutrophils, reciprocally enhancing SC proliferation and muscle repair efficiency. Given that the glycolysis pathway is regulated by the circadian clock, this signaling cascade contributes to variations in muscle repair efficacy depending on the time of day.

Extending our inquiry to encompass a recent study that highlighted a specific subtype of muscle mesenchymal stromal cells (MSCs) subtype exhibiting pro-inflammatory properties post-injury^44^, we examined transcriptomic comparisons across various cell populations including MuSCs, CD31^+^ endothelial cells, CD45^+^ hematopoietic cells, and MSCs in both homeostatic and 1dpi muscles (Supplementary Fig. 10A). The RNA sequencing data from MuSCs derived from this study corroborated our findings, showing a significant upregulation of genes involved in both glycolysis and inflammation response pathways in 1dpi MuSCs versus QSCs (Supplementary Fig. 10B and 10C). The co-activation of these pathways was further substantiated by an observed elevation in related gene expression across all cell types within the injured muscle niche (Supplementary Fig. 10D). Accordingly, activation of glycolysis and inflammatory response, particularly neutrophil migration pathways were consistently observed concomitantly associated with the early hypoxic stage at 1dpi compared to uninjured conditions in all these cell types (Fig. 7B). In addition, a parallel analysis conducted on MSCs from *mdx* mice^44^, a model of Duchenne’s muscular dystrophy, revealed a similar trend in gene expression and signaling pathway activation during spontaneous muscle regeneration (Fig. 7B and Supplementary Fig. 10D). These observations collectively underscore a ubiquitous cellular response to hypoxia that spans across different muscle cell populations.

Taken together, our study delineates a dual role of cytosolic NAD^+^ regeneration during anaerobic glycolysis in hypoxic conditions: it bolsters cell proliferation and concurrently modulates pro-inflammatory signaling, as induced by the activation of PARP1 and NFκB pathways. This process is crucial for the chemotactic recruitment of neutrophils and their interaction with MuSCs, facilitating stem cell activation and proliferation. The circadian clock further influences this process by fine-tuning the hypoxia-induced anaerobic NAD^+^ regeneration, imparting time-of-day-dependent differences in muscle repair efficacy (Fig. 7C).

## Discussion

Metabolic processes are essential for biological functions, with glycolysis being a key player in energy production by converting glucose into pyruvate, which fuels both aerobic and anaerobic pathways^45^. Notably, glycolysis is important for rapidly dividing cells, including tumor and immune cells, to meet their energy demands rapidly. Immune cells predominantly rely on glycolysis when activated, to bolster an inflammatory response, but switch back to oxidative phosphorylation (OXPHOS) after the inflammation subsides^46^. This metabolic shift is closely associated with inflammatory signaling, highlighted by increased glycolysis driving inflammatory markers in endothelial cells via HIF1α upregulation^47^, and the dampening of NF-κB activation when glycolysis is inhibited^48,49^. Similarly, our findings reveal elevated NF-κB signaling (Supplementary Fig.5) and inflammatory marker expression (Fig. 3C) in myoblasts under hypoxic conditions, aligning with known glycolysis-inflammation dynamics. Mechanistically, glycolytic intermediates and enzymes have been implicated in modulating inflammation. For example, the accumulation of phosphoenolpyruvate (PEP) in T cells encourages a pro-inflammatory phenotype^50,51^, and lactate boosts pro-inflammatory activities through NF-κB pathways^52,53^. In addition, certain glycolytic enzymes act as post-transcriptional regulators of inflammatory genes by binding to their mRNAs^54,55^. These findings underscore the significant impact of glycolysis on immune cell activity.

The reason why rapidly proliferating cells discard carbon by expelling pyruvate-converted lactate remains unclear, though recent insights suggest a preference for converting pyruvate to lactate via LDH when the need for NAD^+^ regeneration surpasses ATP demand^56^. While glycolysis facilitates NAD^+^ regeneration critical for maintaining its flux and ATP production, our study unveils added complexity within SCs during early muscle repair. Like cancer cells, glycolysis dominates in proliferating myoblasts even under highly oxygenated culture conditions^57^, underscoring its importance in supporting SC proliferation. Nonetheless, given that quiescent SCs take more than 24 hours to commence and complete their first division^58^, while the rate of glycolysis in SCs is reported to increase as soon as they break quiescence, reaching threefold within 3 h of activation^59^, ATP might not be their primary need immediately post-injury. Our findings indicate that beyond supporting glycolysis, NAD^+^ regeneration is key for immune modulation, activating PARP and NF-κB pathways to induce chemokine production, thus orchestrating neutrophil recruitment and SC interaction. This mechanism is backed by the exclusive induction of *Parp1* (Fig. 3K) and enhanced expression of genes associated with DNA damage responses, a known trigger for PARP1 activation^60^ (Fig. 1E) in 1dpi ASC versus QSCs. Given the rapid NAD^+^-consuming capacity of PARP1^61^, and its ability to prioritize NAD^+^ over Sirtuin proteins^62^, our results suggest that alongside sustaining glycolytic flux, NAD^+^ regeneration through PARP1 activation is crucial for immune modulation in the initial muscle regeneration phase. Overall, these findings expand our understanding of the metabolic-immune interface, indicating that metabolic pathways contribute to immune responses beyond energy provision.

Neutrophils play an indispensable role in the innate immune system, serving as the frontline defense against pathogens^63^. They navigate from the bone marrow to infection or inflammation sites by adhering to the leukocyte adhesion cascade^64^. Interestingly, this migration exhibits a circadian rhythm, potentially linked to the cyclic expression of genes related to migration, inflammation, and survival, governed by the neutrophils’ intrinsic circadian clock^65^. Our findings demonstrate that NAD^+^ regeneration, triggering chemokine release, significantly contributes to neutrophil recruitment across different muscle cell types. Furthermore, the circadian clock intricately modulates glycolytic activity by controlling the HIF1α-mediated response to hypoxia^30,66^. In particular, our previous research highlighted that *Bmal1* in SCs or myoblasts regulates the expression of critical glycolytic genes, including *Glut1*, *Hk2*, and *Ldha*, influencing glycolysis and the anaerobic regeneration of NAD^+^ ^4^. Thus, the circadian regulation of neutrophil infiltration is also influenced by peripheral clock residing in the surrounding cell populations.

Muscle injury leads to the necrosis and degeneration of muscle fibers, accompanied by hematoma formation. Neutrophils rapidly migrate to the injury site within hours of damage^67^. Studies involving mice, where intraperitoneal injections of antisera targeting neutrophils were used alongside myotoxin injections, highlight the critical role of neutrophils in efficient muscle repair^68^. Neutrophils could contribute to muscle regeneration by clearing tissue debris^69^ and by recruiting monocytes and macrophages through the release of inflammatory cytokines and chemokines^70^. A recent *in vitro* study further underscored the influence of neutrophils on muscle regeneration. When myoblasts were treated with media conditioned by activated neutrophils, there was a notable promotion of myoblast proliferation and a suppression of differentiation compared to treatment with basal media^32^. Specifically, the study found that inhibiting the neutrophil-derived factor Ccl3 in the conditioned media nullified the proliferative effects, while blocking Mmp-9 encouraged myoblast differentiation^32^. Intriguingly, *Ccl3* expression was significantly higher at ZT16 compared to ZT4 in the Neutrophil_1 population on day one post-injury (data not shown), suggesting that the interaction between SCs and neutrophils leads to TOD-dependent variations in SC proliferation. Further studies are needed to understand whether TOD-dependent regulation of neutrophil activity can reciprocally impact SC proliferation and myogenic capacity.

Within the subset of neutrophil chemotaxis-related genes that were upregulated in ASCs at 1dpi compared to QSCs, the expression of *Ccl2* showed TOD-dependent variations, being significantly higher in ASCs at ZT16 than at ZT4 (Fig. 4G and 3A). *Ccl2* was the only chemokine gene among the top ligands that exhibited high activity in inducing target gene expression in Neutrophils_1 (Fig. 4F and Supplementary Fig. S7). CCL2 is recognized as a potent chemoattractant for neutrophils^19^, and previous study has indicated its capacity to attract substantial neutrophil numbers to the lung even without underlying inflammation^71^. Moreover, experiments in mice deficient in *Ccl2* have revealed impaired infiltration of monocyte/macrophages, leading to an altered regenerative process that favors adipogenic over myogenic differentiation^72,73^. In addition, a decrease in *Ccl2* expression, resulted from CD8 knockout, has been associated with compromised muscle regeneration and increased fibrosis^74^. However, these studies did not explore the role of *Ccl2* expressed by SC during muscle regeneration. Our findings, through transplantation of *Ccl2*-overexpressing SC-derived myoblasts, demonstrated a significant increase in the presence of these exogenous progenitors within the proliferating muscle stem cell pool of regenerating muscles at day 3 post-injury compared to control myoblasts (Fig. 5I). This highlights the key role of neutrophil interactions in the activation and proliferation of SCs.

The circadian regulation of muscle regeneration has been independently established in prior studies from our group and others^4,5^. While the TOD-dependent differences in muscle repair with cardiotoxin are significant, they are not as pronounced as those observed in mice with systemic or SC-conditional core clock gene deletions when assessed histologically at day 7 post-injury^4,5,75^. Moreover, one study noted that the initial differences in regenerating fiber size observed between injuries inflicted at ZT14 and ZT2 seem to equalize by day 14 post-injury^5^. Here, we demonstrate that the effects of TOD of muscle injury can be compounded with repeated injuries. Specifically, repeated injury during the rest period (i.e. ZT4 for a mouse), led to differences in the size of regenerated myofibers, even at one month following the last injury event. Additionally, this coincided with a trend towards changes in muscle function, reinforcing the long-term negative impact of muscle injury during the rest phase, although further studies are required to establish the exact impact of TOD injury on long-term changes in muscle function. Overall, these findings raise important questions regarding the potential pathological aspects of impaired muscle regeneration in conditions such as obesity, diabetes, and aging, where circadian rhythms are continuously disrupted.

In summary, we have revealed a new mechanism that links NAD^+^ regeneration during anaerobic glycolysis with neutrophil modulation. This interaction, in turn, affects the activation and proliferation of muscle stem cells, thereby influencing the effectiveness of muscle regeneration. Further research is needed to elucidate the role of clock-regulated NAD^+^ metabolism in immune modulation, particularly in the context of acute and chronic disease states.

## Acknowledgements

We would like to thank Dr. Joseph Bass, Dr. Grant Barish, Dr. Sudarshan Dayanidhi, and all members of the Peek, Bass and Barish labs for helpful discussions. We thank the Transgenic and Targeted Mutagenesis Laboratory at Northwestern. This research was supported by the NIH NIDDK grants R01DK123358 and P30DK020595 (C.B.P)., the NIH NIAMS grant K08 AR081391 (I.R.), the NIH NIA grant 5P01AG049665-09 (N.S.C.), and the T32 HL076139-11 (C.R.R.).

## Author Contributions

P.Z. and C.B.P. designed the studies. P.Z. performed and analyzed all experiments unless indicated otherwise. A.V.T., A.W.T.S., and N.R.G. provided technical assistance. Y.Z. provided assistance with bioinformatic analysis of the single-cell sequencing data. I.R, A.W., and R.L.L. performed and analyzed the muscle contractile force experiment. N.S.C. provided LbNOX mice and input on experimental design. C.B.P. and Z.P. wrote the manuscript.

## Supplemental information

Supplementary Figures 1-10. Supplemental Figure Legends and References Cited.

## Methods

### Mice

C57BL/6J mice (#000664) and B6.Cg-Pax7^tm1(cre/ERT2)Gaka^/J mice (#017763) were acquired from The Jackson Laboratory. CytoLbNOX LSL/LSL mice, characterized by a Lox-Stop-Lox cassette upstream of CytoLbNOX and an IRES-linked EGFP, were kindly provided by Dr. Navdeep Chandel at Northwestern University. Pax7-zsGreen mice were generously supplied by Dr. Michael Kyba at the University of Minnesota. To achieve satellite cell (SC)-conditional overexpression of CytoLbNOX, we crossed CytoLbNOX LSL/LSL mice with B6.Cg-Pax7^tm1(cre/ERT2)Gaka^/J mice, resulting in CytoLbNOX^fx/fx^Pax7^tm1(cre/ERT2)^ and control CytoLbNOX^+/+^Pax7^tm1(cre/ERT2)^ mice. CytoLbNOX expression was initiated through five consecutive daily intraperitoneal injections of tamoxifen (20 mg/ml in corn oil, 100 mg/kg body weight). The mice were housed under a 12:12 light:dark cycle. All procedures were carried out with male mice aged 8-12 weeks, in strict accordance with the guidelines of the Institutional Animal Care and Use Committee at Northwestern University.

### Neutrophil depletion

Neutrophil depletion *in vivo* was conducted through intraperitoneal injections of 300 μg the 1A8 monoclonal anti-mouse Ly6G antibody, administered daily over a period of three consecutive days. As a control, the corresponding rat IgG2a isotype control antibody was injected in a similar manner. The efficacy of neutrophils and monocyte depletion post-antibody treatment was verified by flow cytometry analysis.

### Skeletal muscle injury

Cardiotoxin-induced tibialis anterior (TA) muscle injury was conducted according to methodologies previously described^4^. Mice underwent anesthesia via intraperitoneal administration of a ketamine (80 mg/kg) and xylazine (10 mg/kg) mixture. Subsequent to the removal of fur from the injection site, TA muscles received a 50 μl injection of a 10 μM cardiotoxin solution in saline, delivered using a 0.5-mL BD insulin syringe.

### Histology

To evaluate muscle regeneration, TA muscles were harvested at indicated time points after injury and enveloped in a thin layer of Tissue-Tek O.C.T. compound. The samples were snap-frozen in liquid nitrogen-chilled isopentane and preserved at -80°C for later processing. Continuous 10-μm thick cross-sections of the frozen TAs were prepared using a Leica CM1860 cryostat. For hematoxylin and eosin (H&E) staining, sections were air-dried overnight at room temperature. Subsequent rehydration in PBS for 5 minutes was followed by fixation in 10% formalin. The nuclei were then stained with hematoxylin for 5 minutes, with excess stain removed by rinsing under tap water for 5 minutes and briefly dipping in clarifier. Eosin was applied for 30 seconds to stain the cytoplasm and connective tissues. The sections were dehydrated through a graded series of ethanol, cleared in xylene, and mounted with Permount^TM^ mounting medium. The prepared slides were examined using a Keyence BZ-X800 microscope. Cross-sectional area (CSA) of myofibers was quantified using FIJI software^76^ with LabelsToROIs plugin^77^.

### Immunohistochemistry

Tissue sections were allowed to air-dry for 30 minutes at room temperature and were then fixed with 4% paraformaldehyde and permeabilized using 0.25% Triton X-100 for 15 minutes. For blocking, sections were treated with phosphate-buffered saline (PBS) containing 5% goat serum, 2% bovine serum albumin (BSA), and 1% Tween-20 for 1 hour. This was followed by an overnight incubation at 4°C with primary antibodies, including anti-Laminin (1:100), anti-Ly-6G (1:100) in a diluent buffer of PBS with 0.5% goat serum, 2% BSA, and 1% Tween-20. After washing in PBS thrice, the sections were incubated with an Alexa Fluor-conjugated secondary antibodies at a 1:1000 dilution for 1 hour at room temperature. Sections were then mounted using VectaShield HardSet antifade mounting medium with DAPI for visualization under a Keyence BZ-X800 microscope.

### Immunofluorescence

Primary myoblasts were rinsed with ice-cold PBS and fixed with methanal for 15 minutes at –20°C. They were then permeabilized using 0.2% Triton X-100 and blocked to prevent non-specific binding with 1% bovine serum albumin (BSA) for 1 hour at room temperature. This was followed by incubation with an anti-Poly(ADP-ribose) polyclonal antibody at a 1:500 dilution overnight at 4°C. The following day, the myoblasts were washed thrice with PBS and incubated with an Alexa Fluor 594-conjugated goat anti-rabbit IgG secondary antibody at a 1:1000 dilution in 1% BSA for 1 hour at room temperature. Nuclear staining was performed with DAPI (5 μg/mL in PBS) ffor 5 minutes, and the samples were visualized under an Evos FL cell-imaging microscope.

### *In vivo* muscle contractility measurement

Mice were anesthetized using a precision vaporizer (Somnoflo; Kent Scientific) with initial induction of 2.5% isoflurane. After induction, the mouse was transferred to a 37°C motor stage (Aurora Scientific) and the appropriate leg was shaved from ankle to hip.

Dorsiflexion torque was measured as previously described^78^ With the foot fixed to the footplate attached to a 300C series dual mode lever. Briefly, two needle electrodes were placed subcutaneously on either side of the peroneal nerve to activate dorsiflexors. Then, a torque-frequency curve was generated from 10 Hz to 150 Hz as previously described^78^. Torque was recorded at each frequency. ZT4 mice (injured and uninjured) were tested at 9 am while ZT16 mice (injured/uninjured) were tested at 1 pm.

### Retrovirus production

The cDNA open reading frame (ORF) for mouse chemokine (C-C motif) ligand 2 (Ccl2) was ordered from GenScript and subsequently amplified by PCR using Phusion^TM^ High-Fidelity DNA Polymerase & dNTP Mix. The amplified sequence was cloned between the Xho I and EcoR I restriction sites of the retroviral mammalian expression vector MIGR1^79^. To produce retrovirus carrying the Ccl2 gene, 293T cells were transfected with a DNA comprising 1.5 μg of pCL-Eco^80^ and 2.5 μg MIGR1-Ccl2, or MIGR1 vector using Lipofectamine^TM^ 3000 Transfection Reagent according to the provided instructions. The retrovirus-enriched supernatant was collected 24 hours post-transfection and was then filtered through a syringe filter with a pore size of 0.45 μm to ensure purity.

### Muscle cell isolation

For the *in vitro* culture of myoblasts, muscle cells were extracted from the entire hindlimb muscles. Bulk RNA-sequencing involved the use of Pax7-zsgreen mice, which were sequentially perfused with 50 ml of ice-cold PBS, followed by 50 ml mild fixative (0.5% paraformaldehyde), and subsequently quenched with 50 ml 2M glycine, as previously reported^36,81^. SCs were collected from the TA muscles on both sides. For single-cell RNA-sequencing, mononucleated muscle cells were isolated from TA muscles on both hindlimb. Muscle cell preparation was performed by following previously established procedures^4^. Specifically, excised muscle tissues were digested in Hams F-10 medium supplemented with 5 mg/mL collagenase D and 5 mg/mL dispase II for 30 min at 37°C with shaking at 500 rpm. The enzymatic digestion was terminated by adding a fivefold volume of ice-cold cell suspension buffer (Ham’s F-10 with 10% FBS, 3 mM EDTA). The digested muscle tissue was then subjected to mechanical dissociation through vigorous pipetting (20 times with a 10-ml pipette) to release MuSCs from the myofibers. The cell suspension was filtered through 70- and 40-μm nylon mesh strainers in sequence, and the filtrate was collected in a 15-ml conical tube and centrifuged at 500×g for 5 minutes. The resulting cell pellets were resuspended in 1 mL of Pre-Sort buffer, with Propidium Iodide serving as an indicator of cell viability. For bulk RNA-sequencing, zsgreen-positive mononuclear SCs were sorted using a BD FACSAria II cell sorter.

Myoblasts preparation was performed with slight modification to a previously established preplating technique^82^. Initially, the crude suspension was seeded and expanded on a 60-mm dish coated with Matrigel, in myoblast growth medium consisting of Hams F-10 with 20% fetal bovine serum (FBS), 1% penicillin–streptomycin (PS), and 2.5 ng/ml basic fibroblast growth factor (bFGF) for three days. Following this, cells underwent trypsinization and were subjected to a preplating process on a regular tissue culture-treated 60-mm dish for 1 hour, before being transferred to another Matrigel-coated dish. This procedure was repeated three times to significantly reduce the presence of nonmyoblast cells. The resultant myoblast-enriched cultures were then maintained in a growth medium composed of DMEM/Hams F-10 (1:1) with 20% FBS, 1% PS, and 2.5 ng/mL bFGF. The purity of myoblast, verified to be over 95% using immunofluorescence with a Pax7-antibody (Developmental Studies Hybridoma Bank), underscored the effectiveness of this method. Induction of Cyto-LbNOX was conducted no sooner than 48 hours after primary myoblasts were infected with either an adenovirus expressing Cre recombinase (Adv-Cre) or a control empty vector (Adv-Ctrl) from Vector Biolabs.

### SC Transplantation

SCs from wildtype C57BL/6 mice were isolated based on a combination of cell surface markers CD45^-^CD31^-^Sca1^-^VcamI^+ 42^. These cells were then transduced overnight with either a control or *Ccl2*-expressing retrovirus, in the presence of 8 µg/ml polybrene within wells coated with Matrigel. To prepare for transplantation, TA muscles in recipient wildtype C57BL/6 mice underwent injury preconditioning with a 50 solution of 10 μM cardiotoxin solution in saline, administered 6 hours prior to cell transplantation. Subsequently, 10,000 SCs, transduced with either control or *Ccl2*-expressing retrovirus as denoted by GFP expression, were sorted, and intramuscularly injected into the pre-injured TA muscles on each side. The cells were delivered in a small volume of 15 μl using 0.3 ml insulin syringes to ensure precise administration.

### Bulk RNA-sequencing

Total RNA was extracted from *in vivo* light-fixed Pax7-zsgreen^+^ SCs using the miRNeasy FFPE Kit, adhering to the supplied instructions with a crucial modification that the digestion of RNA by proteinase K was performed at 56°C for 1 hour as previously reported^36^. For the initial double-stranded cDNA synthesis, amounts ranging from 10 picograms to 10 nanograms of total RNA were employed using the SMART-seq v4 Ultra Low Input RNA Kit from Takara Bio, following the protocol included within the kit. After PCR amplification, the resulting cDNA was purified using the AMPure XP Beads. This purified cDNA underwent assessment using the Agilent 2100 Bioanalyzer paired with Agilent’s High-Sensitivity DNA Kit. Three hundred picograms of the full-length cDNA, derived from the SMART-seq v4 Ultra Low-Input RNA Kit, was then prepared for sequencing using the Nextera XT DNA Library Preparation Kit. The PCR-amplified, size-selected cDNA fragments were both visualized and quantified using the Agilent 2100 Bioanalyzer in conjunction with Agilent’s High-Sensitivity DNA Kit, and the Qubit dsDNA HS Assay Kit, respectively. The pooled libraries were sequenced using NovaSeq 6000 SP Reagent Kit v1.5 (100 cycles) on a NovaSeq 6000 system, executing a paired-end run (51 bp, repeated twice) to achieve a sequencing depth of approximately 50 million reads per sample.

### RNA sequencing analysis

Raw BCL files were processed into demultiplexed, paired-end FastQ files using bcl2fastq conversion software (v2.19.1). Subsequently, adaptor sequences were trimmed from these files using Trimmomatic (v0.33)^83^. The processed reads were then aligned to the mm10 Mus musculus reference genome using the STAR aligner (v2.5.2)^84^. Gene alignment counts were determined using RSEM (v1.3.3)^85^. Isoform-level quantifications obtained from RSEM were integrated into DESeq2^86^ for differential expression analysis, facilitated by the tximport package^87^, with settings adjusted to “rsem” and both “txIn” and “txOut” flags set to “TRUE.” The DESeq2 ‘s results function was employed to identify differentially expressed genes (DEGs), which were then subjected to multiple testing correction using the Benjamini-Hochberg method to calculate adjusted P-values. Visualization of gene counts for DEGs was achieved using the pheatmap (v1.0.12) and EnhancedVolcano (version 1.2.0) packages within R (v3.6.1). Finally, KEGG and Elsevier Pathway Collection pathways enrichment as well as gene ontology analyses were conducted using EnrichR^88^, ShinyGO (v0.66)^89^, and Fast Gene Set Enrichment Analysis (doi: https://doi.org/10.1101/060012).

### Single-cell RNA sequencing

Muscle cells were isolated from both intact and cardiotoxin-injured TA muscles on both sides of hindlimb at ZT4 and ZT16 on days 0 (intact), 1, and 3 post-injuries. The initial step involved the elimination of red blood cells using RBC Lysis Buffer, followed by resuspension of the muscle cells in 1 ml of Pre-Sort buffer. Cell viability was assessed with Propidium Iodide. To ensure accurate cell sorting, doublet exclusion was strictly conducted by comparing the height to the area for both forward scatter and side scatter signals. Approximately 100,000 propidium iodide-negative single muscle cells were then sorted from each sample using a BD FACSAria II cell sorter. Subsequent sample processing was completed at the NUSeq Core Facility at Northwestern University. Briefly, the cell number and viability were further evaluated using the Nexcelom Cellometer Auto2000, employing the AOPI fluorescent staining method. For single-cell RNA sequencing, 10,000 cells from each condition were loaded into a Chromium Controller (10X Genomics) using a Chromium Next GEM Chip G (10X Genomics). These were processed to form single-cell gel beads in the emulsion (GEM) following the manufacturer’s instructions. cDNA synthesis and library preparation were performed using the Chromium Next GEM Single Cell 3′ Reagent Kits v3.1 and Single Index Kit T Set A (10X Genomics) according to the provided manual. Quality control for constructed library was performed by Agilent Bioanalyzer High Sensitivity DNA kit (Agilent Technologies) and Qubit DNA HS assay kit for qualitative and quantitative analysis, respectively. The multiplexed libraries were pooled and sequenced using NovaSeq 6000 S1 Reagent Kit v1.5 (100 cycles) on Illumina NovaSeq 6000 sequencer with 2 × 50 paired-end kits using the following read length: 28 bp Read1 for cell barcode and UMI (unique molecular identifier), 8 bp I7 index for sample index and 91 bp Read2 for transcript.

### Single-cell RNA sequencing analysis

For demultiplexing and read counting, Cell Ranger (v1.0.0) was used. The Seurat package (v4.0.0)^90^ in R facilitated data preprocessing and visualization. Quality control measures and cell selection criteria were stringently applied; cells exhibiting less than 200 or more than 6000 feature counts (for cells from intact and 1 dpi injured TA muscles) or exceeding 7500 feature counts (for cells from 3 dpi injured TA muscles) were excluded. Genes detected in fewer than three cells were discarded. To account for low-quality and dying cells typically showing high mitochondrial content, cells were selected based on a threshold where only those with less than 5% of reads mapping to the mitochondrial were retained. Additionally, MALAT1, a highly expressed large non-coding nuclear gene, was excluded from downstream analyses.

To identify potential doublets, a two-tiered approach was adopted as per Germain et al., 2021^91^: initially, scDblFinder (v1.4.0) predicted doublets using default settings, followed by manual identification based on the expression of marker genes from different cell types. After refining the dataset to remove low-quality cells, normalization of single cells from all 6 samples was carried out using NormalizeData function, and samples were integrated using a canonical correlation analysis (CCA) based anchor method. The dataset was further adjusted by regressing out the difference between G2M and S phase scores, effectively preserving distinctions between non-cycling and cycling cells while ignoring variations among proliferating cells. Clustering was performed using the first 50 principal components (PC), derived from the top 2,000 most variable features, in a K-nearest neighbor (KNN) graph with a resolution at 0.5. Subsequently, Uniform Manifold Approximation and Projection (UMAP) provided non-linear dimensional reduction and visualization of clusters. Differentially expressed features across clusters were identified using FindAllMarkers function. Cell identities within each cluster were ascribed based on canonical marker genes for skeletal muscle cells, as described in previous studies^14,15^, and markers from the CellMarker 2.0 database^13^. Differential gene expression analyses between clusters utilized the FindMarkers function, setting a threshold of 0.25 for log fold change and 0.05 for the adjusted p-value. Cell lineage and pseudotime inference analysis within MuSC subclusters were conducted using the Slingshot package^38^, with downstream gene expression along trajectories analyzed via the tradeSeq package^39^. Intercellular interaction analysis and visualization among the two neutrophil and MuSC populations at various days post-injury were performed using the CellChat package^18^ with standard parameters. To examine time-of-day-dependent variations in communication between MuSC and neutrophils, the NicheNet package^31^ was employed, positioning MuSC as the “signal sender” to explore the ligands impacting neutrophils.

### Western blotting

Myoblast whole-cell lysates were prepared using CelLytic^TM^ Mammalian Tissue Lysis Reagent, with the addition of protease inhibitors, to ensure protein integrity. For the preparing total protein lysates from muscle tissue, each snap-frozen TA muscle specimen was placed in 500 μl of CelLytic^TM^ Mammalian Tissue Lysis Reagent supplemented with protease inhibitors and homogenized using a TissueLyser II apparatus from Qiagen to achieve uniform cell disruption. Subsequent to homogenization, protein concentrations were determined using the DC Protein Assay Kit, enabling accurate loading for SDS-PAGE gel electrophoresis. Proteins separated by electrophoresis were then transferred onto 0.45-μm nitrocellulose membranes for immunoblotting. For the detection of specific proteins, the membranes were incubated overnight at 4°C with primary antibodies, including anti-HIF1α (1:1000 dilution), Phospho-NF-κB p65 (Ser536; 1:1000 dilution), NF-κB p65 (1:1000 dilution), PAR/pADPr (1:1000 dilution), and β-Actin (1:10,000 dilution). Following primary antibody incubation, the membranes were washed and then incubated with appropriate horseradish peroxidase (HRP)-conjugated secondary antibodies. The protein bands were visualized using an enhanced chemiluminescence (ECL) detection system, allowing for the quantitative analysis of protein expression levels.

### RNA extraction and Real-Time quantitative PCR (qPCR)

For the qPCR analysis, total RNA was isolated from myoblasts using TRIzol^TM^ Reagent, adhering strictly to the guidelines provided by the manufacturer. The quality and quantity of the extracted total RNA were assessed using the NanoDrop^TM^ 2000 Spectrophotometers. RNA samples displaying an OD260/280 ratio greater than 1.80 were deemed suitable and thus advanced to the first strand cDNA synthesis using Applied Biosystems^TM^ High-Capacity cDNA Reverse Transcription Kit, following the instructions detailed in the kit’s protocol. The subsequent Real-Time quantitative PCR was performed using iTaq Universal SYBR Green Supermix from Bio-Rad on a CFX Opus 384 Real-Time PCR System.

### ELISA

The concentration of CCL2 released by myoblasts into the culture medium was quantified using the Mouse MCP-1 ProQuantum Immunoassay Kit following the instructions provided by the manufacturer.

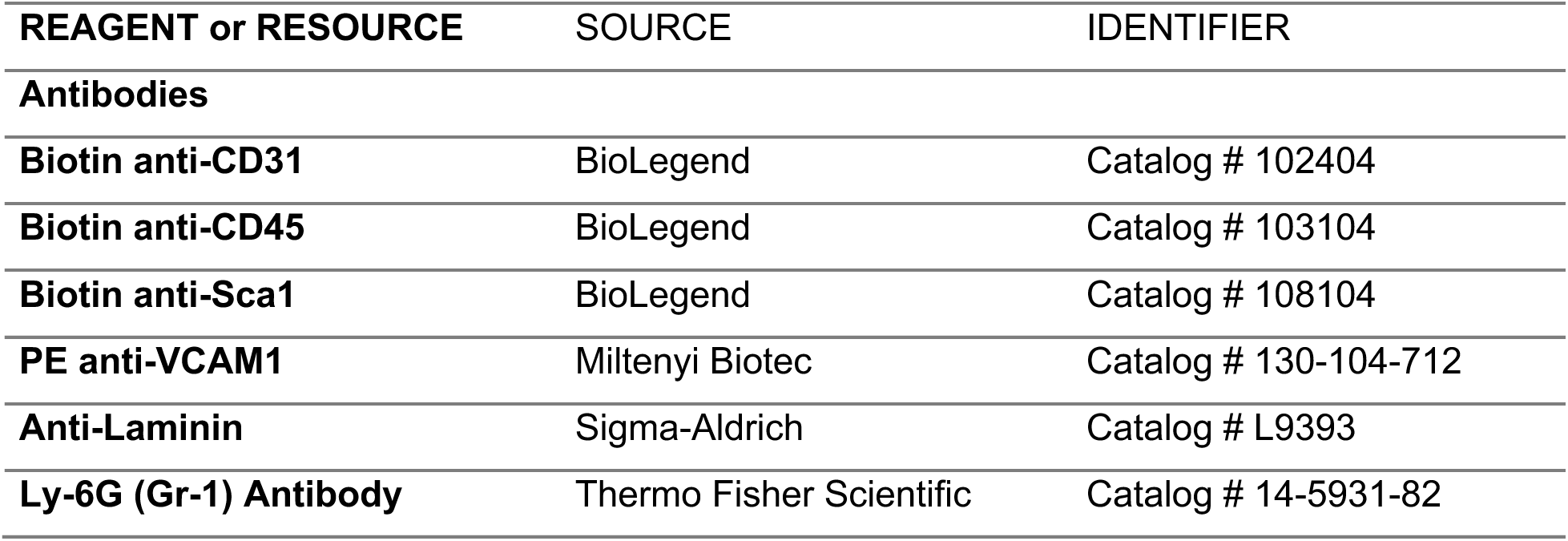
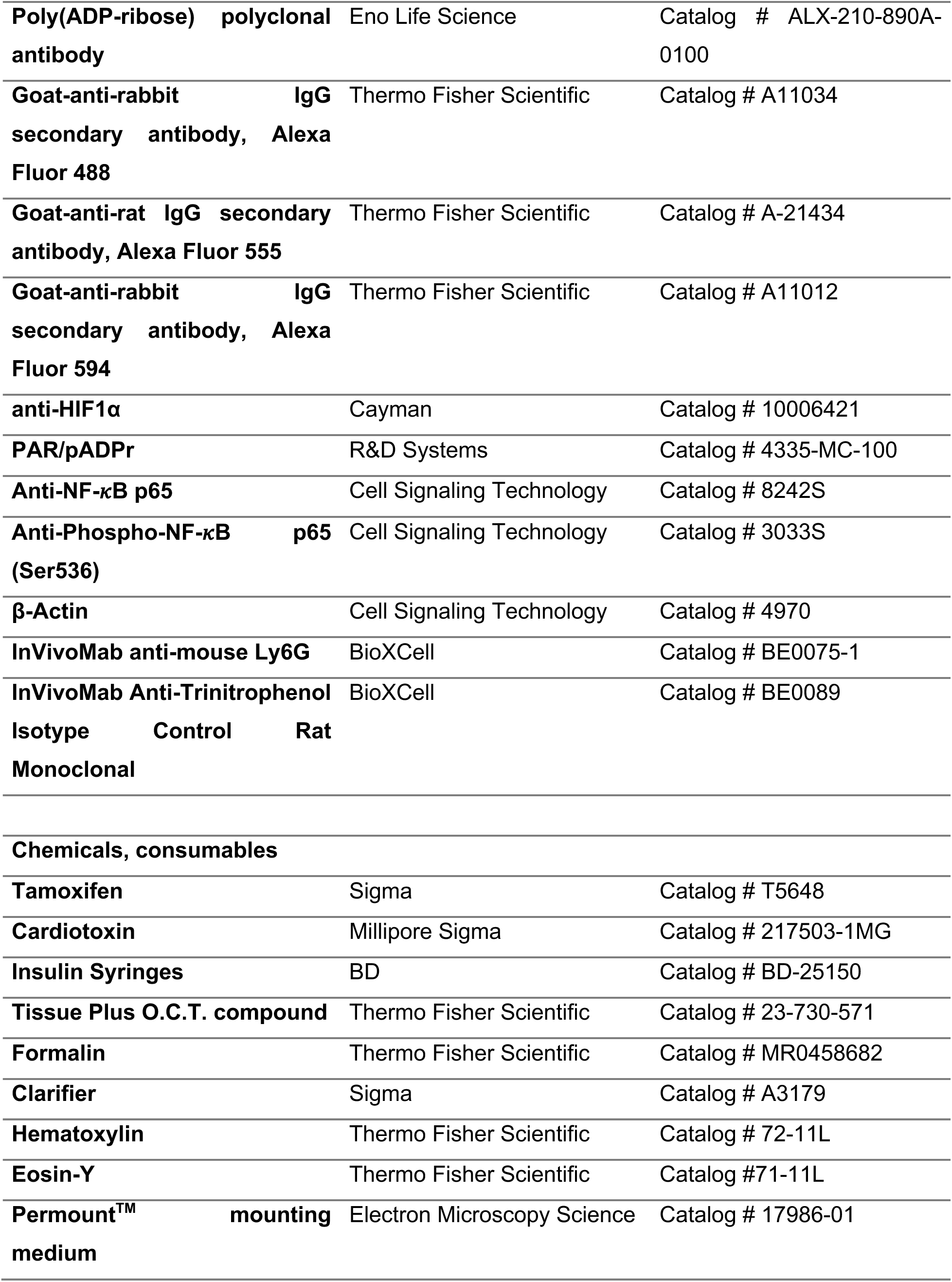
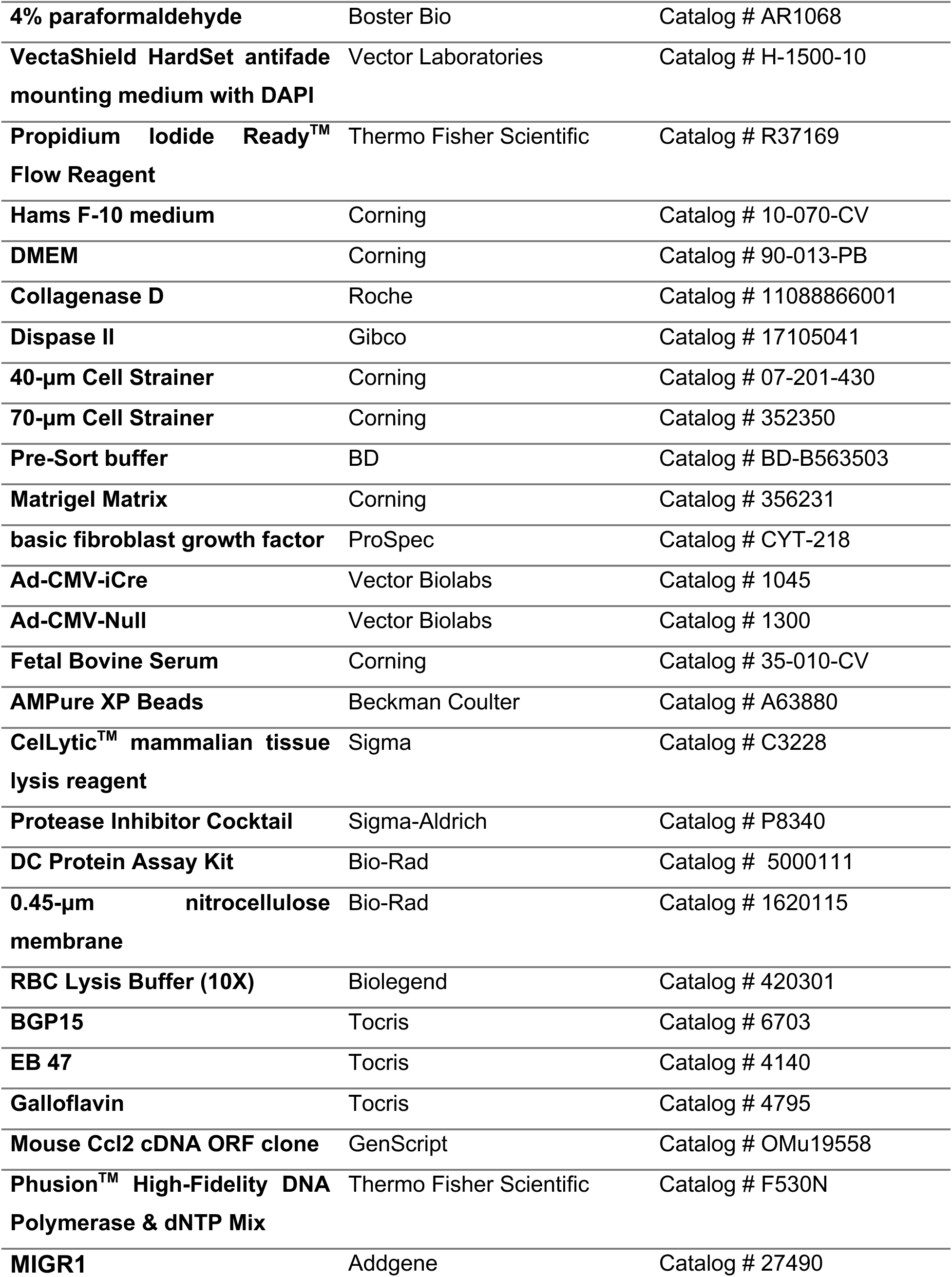
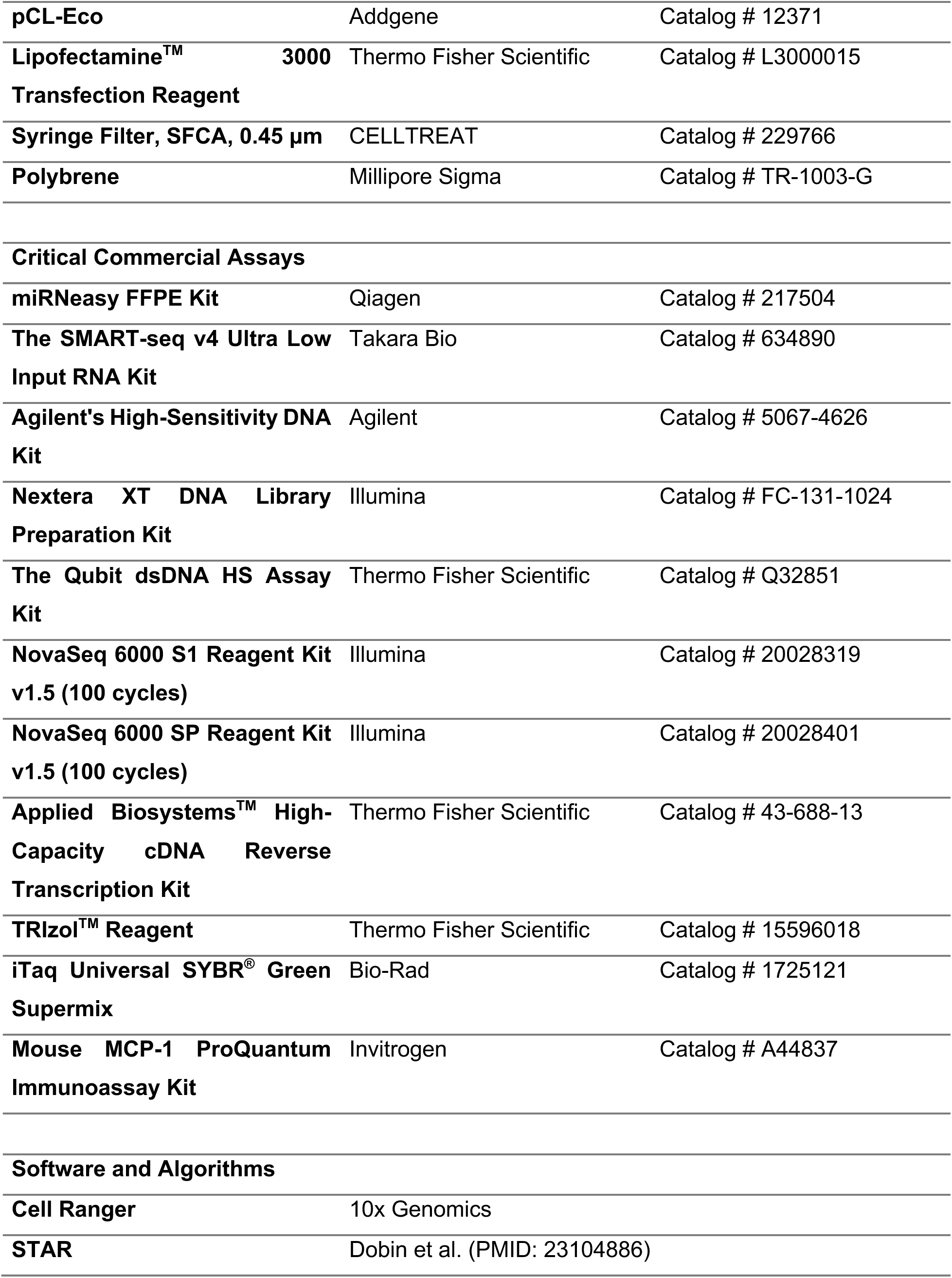
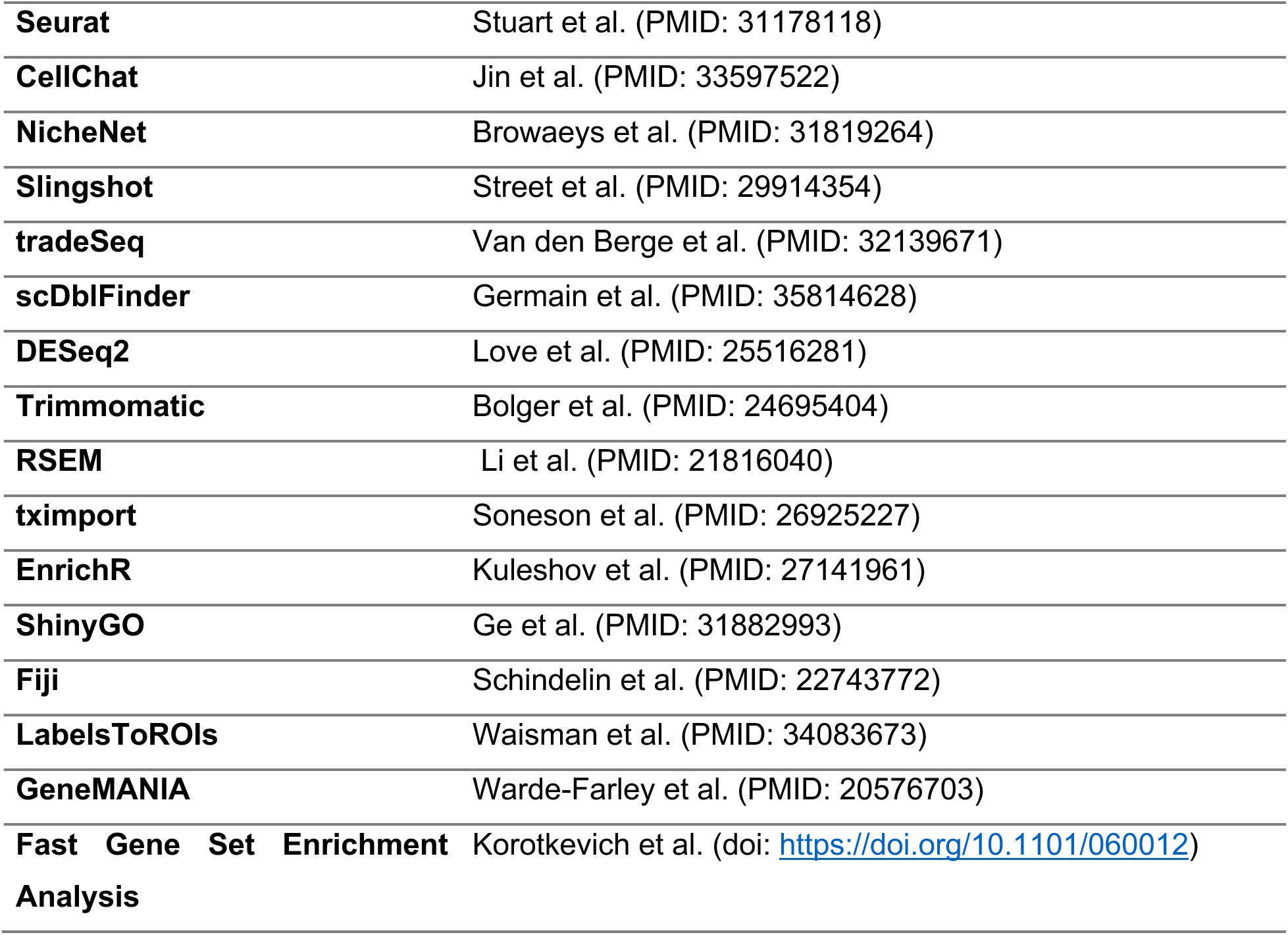

