## Supplemental Materials for "Immunomodulatory Role of the Stem Cell Circadian Clock in Muscle Repair"

#### Figure S1

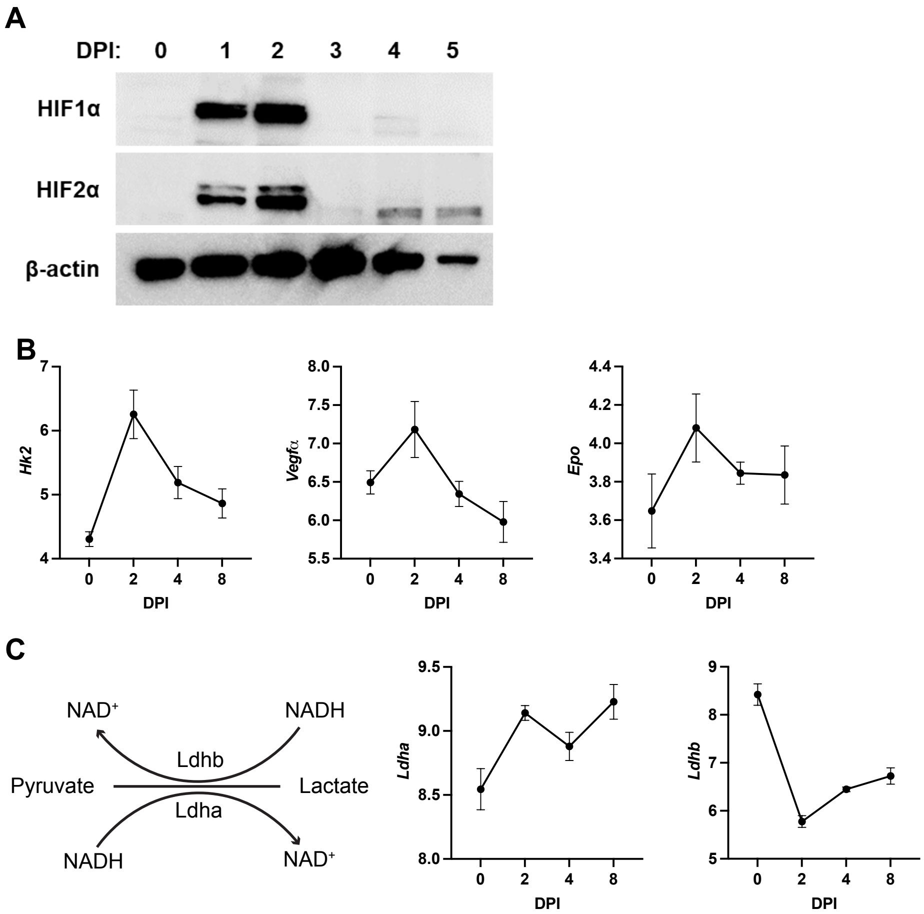
A
DPI: 0 1 2 3 4 5
HIF1α
HIF2α
β-actin
B
Hk2
Vegfα
Epo
DPI
C
NAD⁺
NADH
Ldhb
Pyruvate
Lactate
Ldha
NADH
NAD⁺
Ldha
Ldhb
DPI

### Figure S2

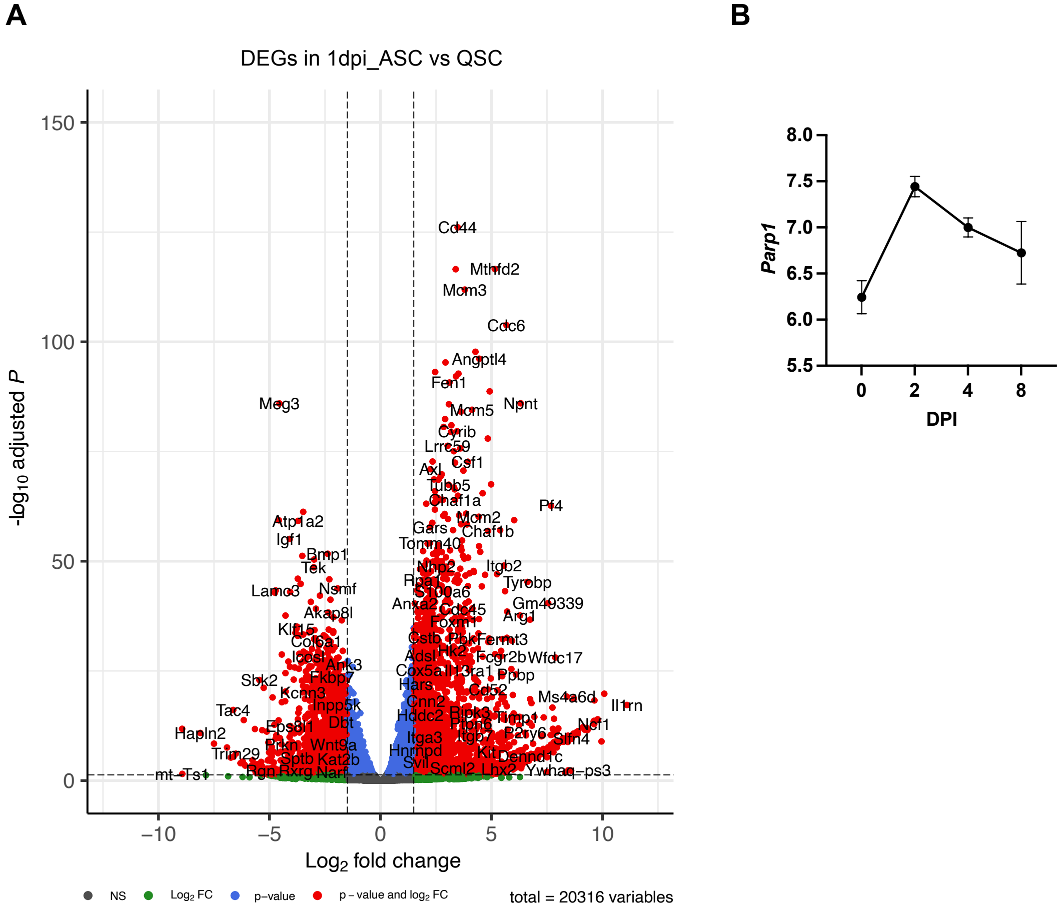
A
B
DEGs in 1dpi_ASC vs QSC
-log10 adjusted P
Log2 fold change
150
100
50
0
−10
−5
0
5
10
NS
Log2 FC
p−value
p − value and log2 FC
total = 20316 variables
Cd44
Mthfd2
Mcm3
Cdc6
Angptl4
Fen1
Meg3
Npnt
Mcm5
Pf4
Atp1a2
Igf1
Tac4
Hapln2
Il1rn
Parp1
DPI
8.0
7.5
7.0
6.5
6.0
5.5
0
2
4
8

### Figure S3

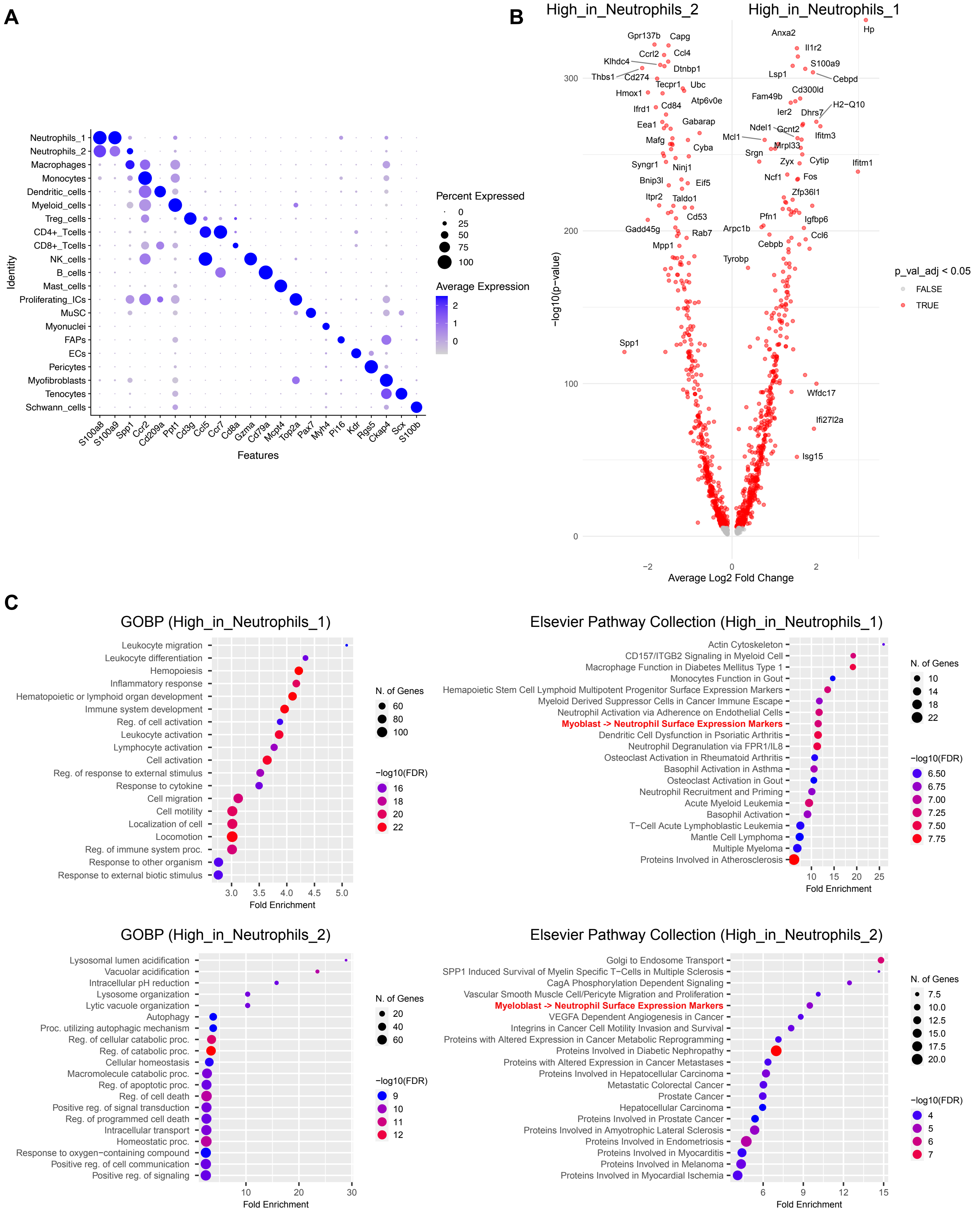
A
B
C
Percent Expressed
Average Expression
Identity
Features
High_in_Neutrophils_2
High_in_Neutrophils_1
-log10(p-value)
Average Log2 Fold Change
p_val_adj < 0.05
FALSE
TRUE
GOBP (High_in_Neutrophils_1)
Elsevier Pathway Collection (High_in_Neutrophils_1)
GOBP (High_in_Neutrophils_2)
Elsevier Pathway Collection (High_in_Neutrophils_2)
N. of Genes
-log10(FDR)
Fold Enrichment
Leukocyte migration
Leukocyte differentiation
Hemopoiesis
Inflammatory response
Hematopoietic or lymphoid organ development
Immune system development
Reg. of cell activation
Leukocyte activation
Lymphocyte activation
Cell activation
Reg. of response to external stimulus
Response to cytokine
Cell migration
Cell motility
Localization of cell
Locomotion
Reg. of immune system proc.
Response to other organism
Response to external biotic stimulus
Actin Cytoskeleton
CD157/ITGB2 Signaling in Myeloid Cell
Macrophage Function in Diabetes Mellitus Type 1
Monocytes Function in Gout
Hemapoietic Stem Cell Lymphoid Multipotent Progenitor Surface Expression Markers
Myeloid Derived Suppressor Cells in Cancer Immune Escape
Neutrophil Activation via Adherence on Endothelial Cells
Myoblast -> Neutrophil Surface Expression Markers
Dendritic Cell Dysfunction in Psoriatic Arthritis
Neutrophil Degranulation via FPR1/IL8
Osteoclast Activation in Rheumatoid Arthritis
Basophil Activation in Asthma
Osteoclast Activation in Gout
Neutrophil Recruitment and Priming
Acute Myeloid Leukemia
Basophil Activation
T-Cell Acute Lymphoblastic Leukemia
Mantle Cell Lymphoma
Multiple Myeloma
Proteins Involved in Atherosclerosis
Lysosomal lumen acidification
Vacuolar acidification
Intracellular pH reduction
Lysosome organization
Lytic vacuole organization
Autophagy
Proc. utilizing autophagic mechanism
Reg. of cellular catabolic proc.
Reg. of catabolic proc.
Cellular homeostasis
Macromolecule catabolic proc.
Reg. of apoptotic proc.
Reg. of cell death
Positive reg. of signal transduction
Reg. of programmed cell death
Intracellular transport
Homeostatic proc.
Response to oxygen-containing compound
Positive reg. of cell communication
Positive reg. of signaling
Golgi to Endosome Transport
SPP1 Induced Survival of Myelin Specific T-Cells in Multiple Sclerosis
CagA Phosphorylation Dependent Signaling
Vascular Smooth Muscle Cell/Pericyte Migration and Proliferation
Myeloblast -> Neutrophil Surface Expression Markers
VEGFA Dependent Angiogenesis in Cancer
Integrins in Cancer Cell Motility Invasion and Survival
Proteins with Altered Expression in Cancer Metabolic Reprogramming
Proteins Involved in Diabetic Nephropathy
Proteins with Altered Expression in Cancer Metastases
Proteins Involved in Hepatocellular Carcinoma
Metastatic Colorectal Cancer
Prostate Cancer
Hepatocellular Carcinoma
Proteins Involved in Prostate Cancer
Proteins Involved in Amyotrophic Lateral Sclerosis
Proteins Involved in Endometriosis
Proteins Involved in Myocarditis
Proteins Involved in Melanoma
Proteins Involved in Myocardial Ischemia

**Figure S4**

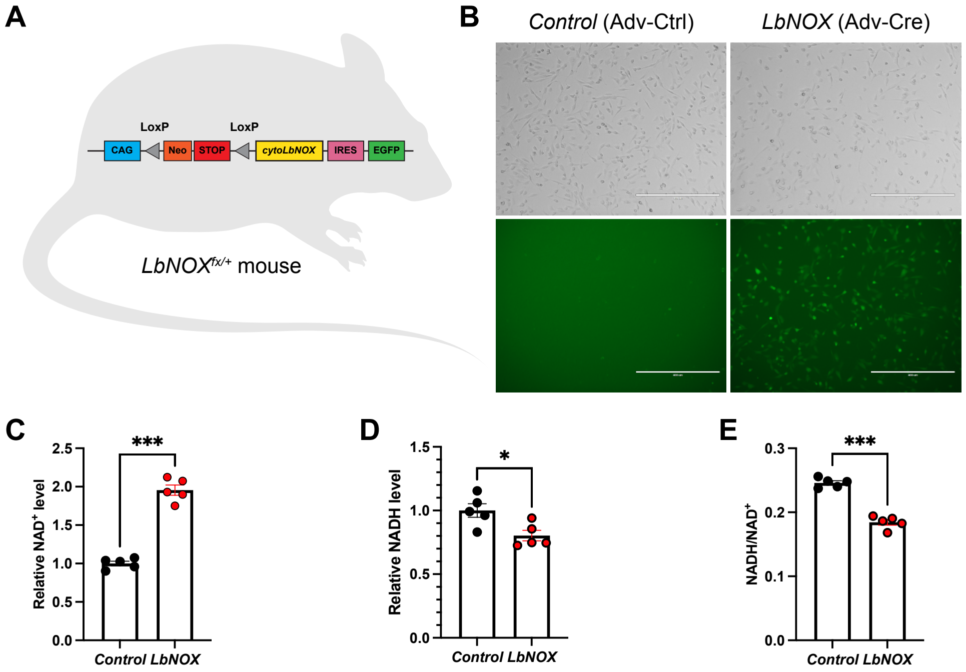

A
LoxP
LoxP
CAG
Neo
STOP
cytoLbNOX
IRES
EGFP
LbNOXfx/+ mouse
B
Control (Adv-Ctrl)
LbNOX (Adv-Cre)
C
***
Relative NAD+ level
2.5
2.0
1.5
1.0
0.5
0.0
Control LbNOX
D
*
Relative NADH level
1.5
1.0
0.5
0.0
Control LbNOX
E
***
NADH/NAD+
0.3
0.2
0.1
0.0
Control LbNOX

#### Figure S5

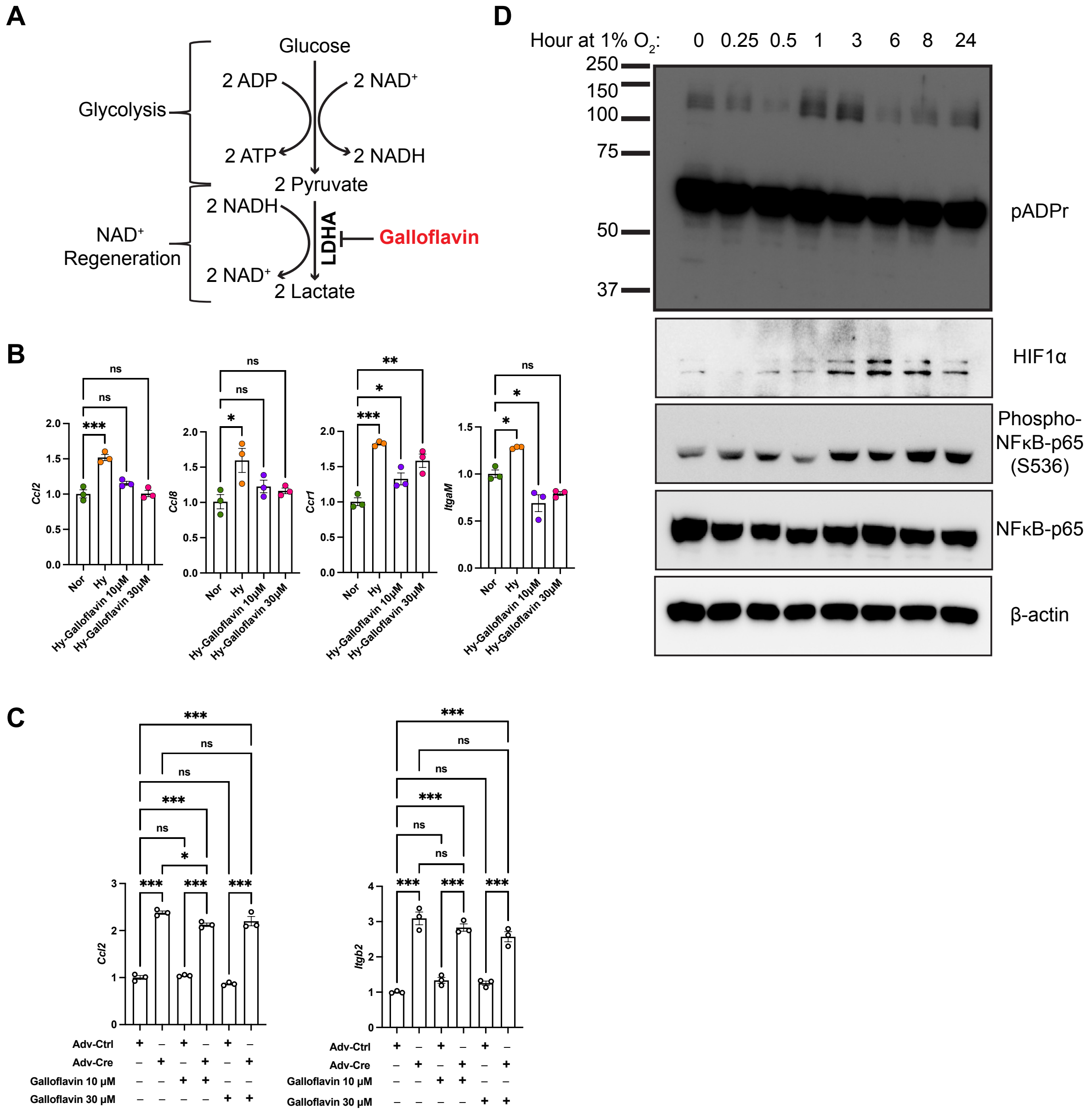

A
Glucose
2 ADP
2 NAD+
Glycolysis
2 ATP
2 NADH
2 Pyruvate
2 NADH
NAD+ Regeneration
LDHA
Galloflavin
2 NAD+
2 Lactate
B
Ccl2
Ccl8
Ccr1
ItgaM
Nor
Hy
Hy-Galloflavin 10μM
Hy-Galloflavin 30μM
C
Ccl2
Itgb2
Adv-Ctrl
Adv-Cre
Galloflavin 10 μM
Galloflavin 30 μM
D
Hour at 1% O2: 0 0.25 0.5 1 3 6 8 24
250
150
100
75
50
37
pADPr
HIF1α
Phospho-NFκB-p65 (S536)
NFκB-p65
β-actin

### Figure S6

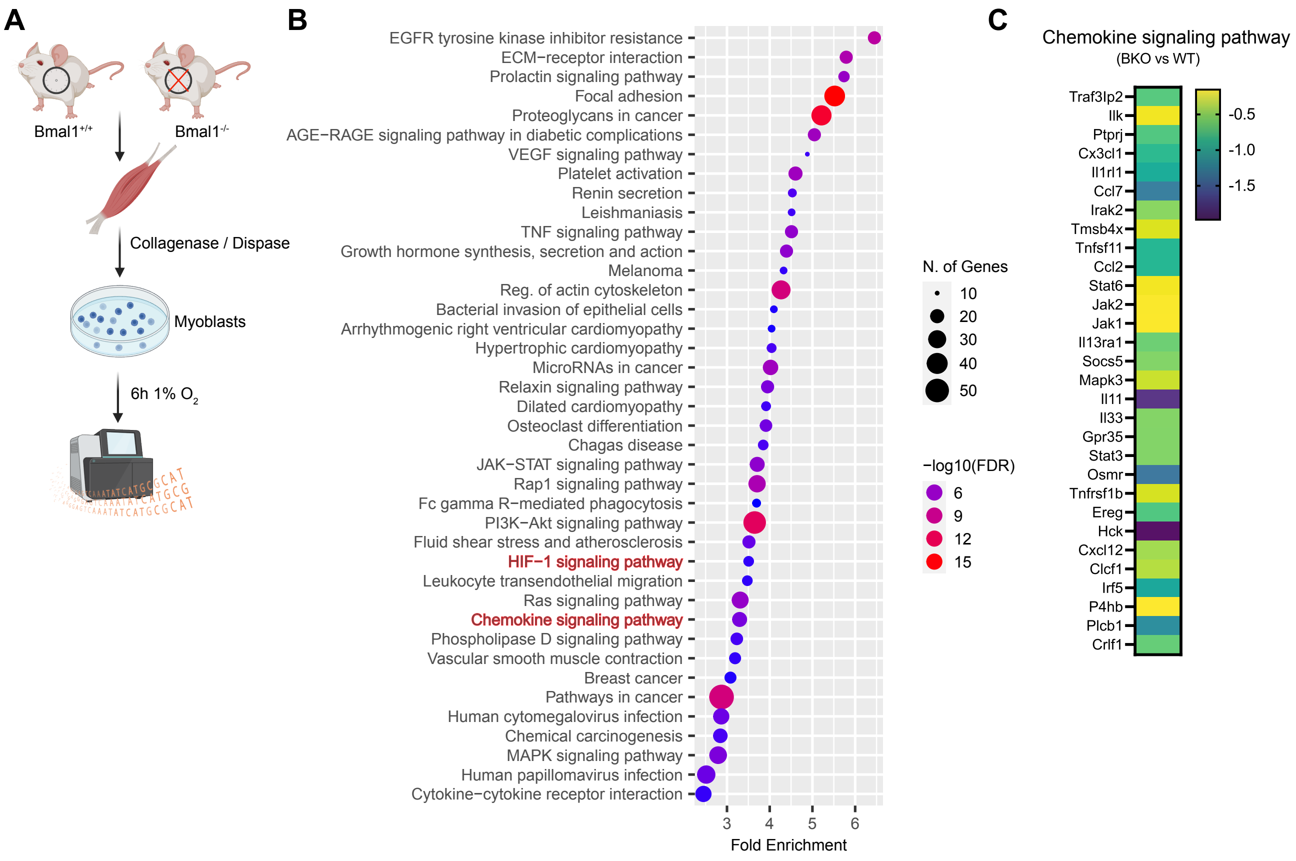

A
Bmal1+/+
Bmal1-/-
Collagenase / Dispase
Myoblasts
6h 1% O2
B
EGFR tyrosine kinase inhibitor resistance
ECM−receptor interaction
Prolactin signaling pathway
Focal adhesion
Proteoglycans in cancer
AGE−RAGE signaling pathway in diabetic complications
VEGF signaling pathway
Platelet activation
Renin secretion
Leishmaniasis
TNF signaling pathway
Growth hormone synthesis, secretion and action
Melanoma
Reg. of actin cytoskeleton
Bacterial invasion of epithelial cells
Arrhythmogenic right ventricular cardiomyopathy
Hypertrophic cardiomyopathy
MicroRNAs in cancer
Relaxin signaling pathway
Dilated cardiomyopathy
Osteoclast differentiation
Chagas disease
JAK−STAT signaling pathway
Rap1 signaling pathway
Fc gamma R−mediated phagocytosis
PI3K−Akt signaling pathway
Fluid shear stress and atherosclerosis
HIF−1 signaling pathway
Leukocyte transendothelial migration
Ras signaling pathway
Chemokine signaling pathway
Phospholipase D signaling pathway
Vascular smooth muscle contraction
Breast cancer
Pathways in cancer
Human cytomegalovirus infection
Chemical carcinogenesis
MAPK signaling pathway
Human papillomavirus infection
Cytokine−cytokine receptor interaction
3
4
5
6
Fold Enrichment
N. of Genes
10
20
30
40
50
−log10(FDR)
6
9
12
15
C
Chemokine signaling pathway
(BKO vs WT)
Traf3Ip2
Ilk
Ptprj
Cx3cl1
Il1rl1
Ccl7
Irak2
Tmsb4x
Tnfsf11
Ccl2
Stat6
Jak2
Jak1
Il13ra1
Socs5
Mapk3
Il11
Il33
Gpr35
Stat3
Osmr
Tnfrsf1b
Ereg
Hck
Cxcl12
Clcf1
Irf5
P4hb
Plcb1
Crlf1
-0.5
-1.0
-1.5

**Figure S7**

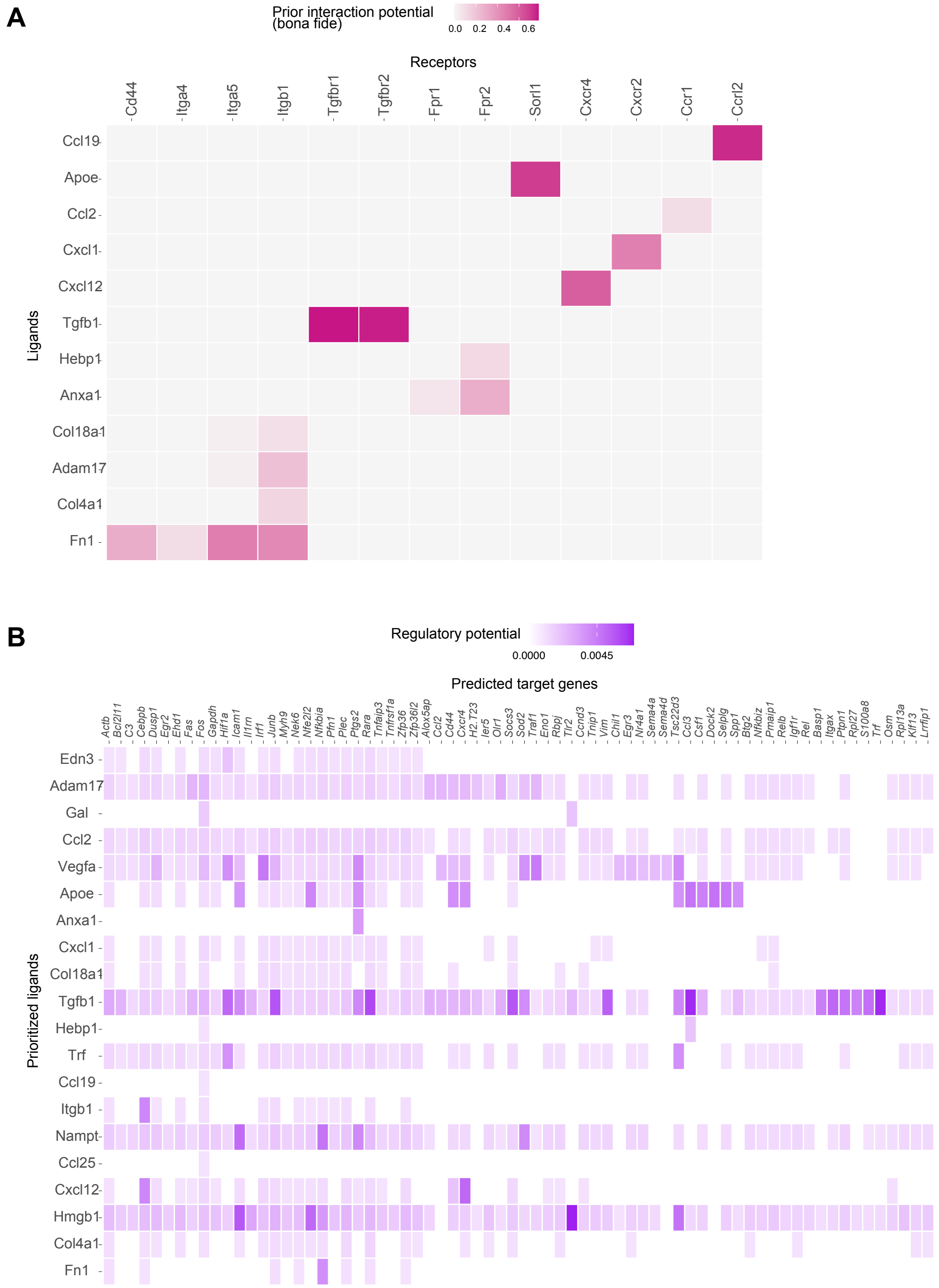
A
Prior interaction potential
(bona fide)
0.0 0.2 0.4 0.6
Receptors
Cd44
Itga4
Itga5
Itgb1
Tgfbr1
Tgfbr2
Fpr1
Fpr2
Sorl1
Cxcr4
Cxcr2
Ccr1
Ccrl2
Ligands
Ccl19
Apoe
Ccl2
Cxcl1
Cxcl12
Tgfb1
Hebp1
Anxa1
Col18a1
Adam17
Col4a1
Fn1
B
Regulatory potential
0.0000 0.0045
Predicted target genes
Actb Bcl2l11 C3 Cebpb Dusp1 Egr2 Ehd1 Fas Fos Gapdh Hif1a Icam1 Il1rn Irf1 Junb Myh9 Nek6 Nfe2l2 Nfkbia Pfn1 Plec Ptgs2 Rara Tnfaip3 Tnfrsf1a Zfp36 Zfp36l2 Alox5ap Ccl2 Cd44 Cxcr4 H2.T23 Ier5 Olr1 Socs3 Sod2 Traf1 Eno1 Rbpj Tlr2 Ccnd3 Tnip1 Vim Chil1 Egr3 Nr4a1 Sema4a Sema4d Tsc22d3 Ccl3 Csf1 Dock2 Selplg Spp1 Btg2 Nfkbiz Pmaip1 Relb Igf1r Rel Basp1 Itgax Ptpn1 Rpl27 S100a8 Trf Osm Rpl13a Klf13 Lrrfip1
Prioritized ligands
Edn3
Adam17
Gal
Ccl2
Vegfa
Apoe
Anxa1
Cxcl1
Col18a1
Tgfb1
Hebp1
Trf
Ccl19
Itgb1
Nampt
Ccl25
Cxcl12
Hmgb1
Col4a1
Fn1

### Figure S8

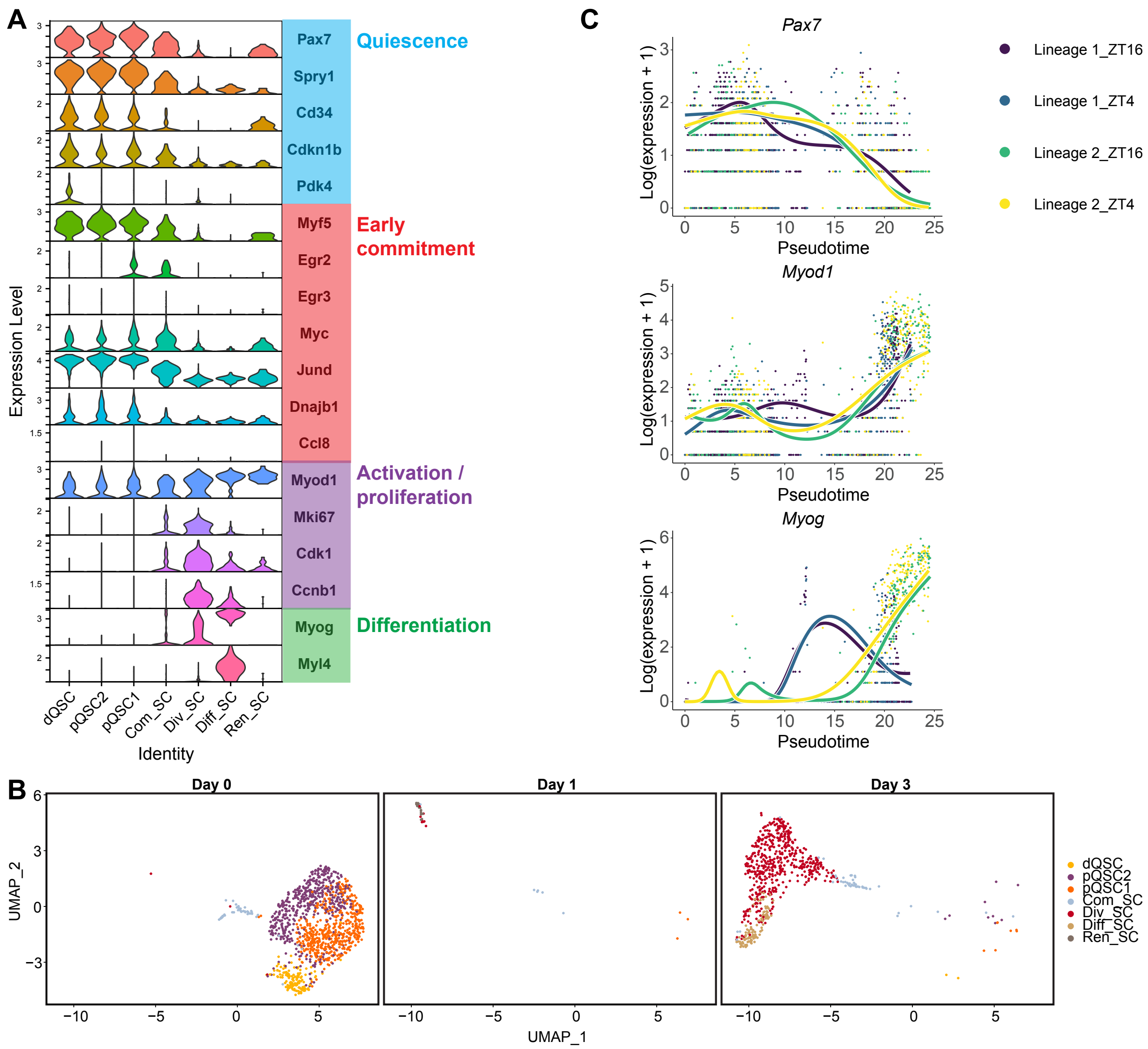
A
Pax7
Spry1
Cd34
Cdkn1b
Pdk4
Quiescence
Myf5
Egr2
Egr3
Myc
Jund
Dnajb1
Ccl8
Early commitment
Myod1
Mki67
Cdk1
Ccnb1
Activation / proliferation
Myog
Myl4
Differentiation
Expression Level
dQSC
pQSC2
pQSC1
Com_SC
Div_SC
Diff_SC
Ren_SC
Identity
B
Day 0
Day 1
Day 3
UMAP_2
UMAP_1
dQSC
pQSC2
pQSC1
Com_SC
Div_SC
Diff_SC
Ren_SC
C
Pax7
Myod1
Myog
Log(expression + 1)
Pseudotime
Lineage 1_ZT16
Lineage 1_ZT4
Lineage 2_ZT16
Lineage 2_ZT4

#### Figure S9

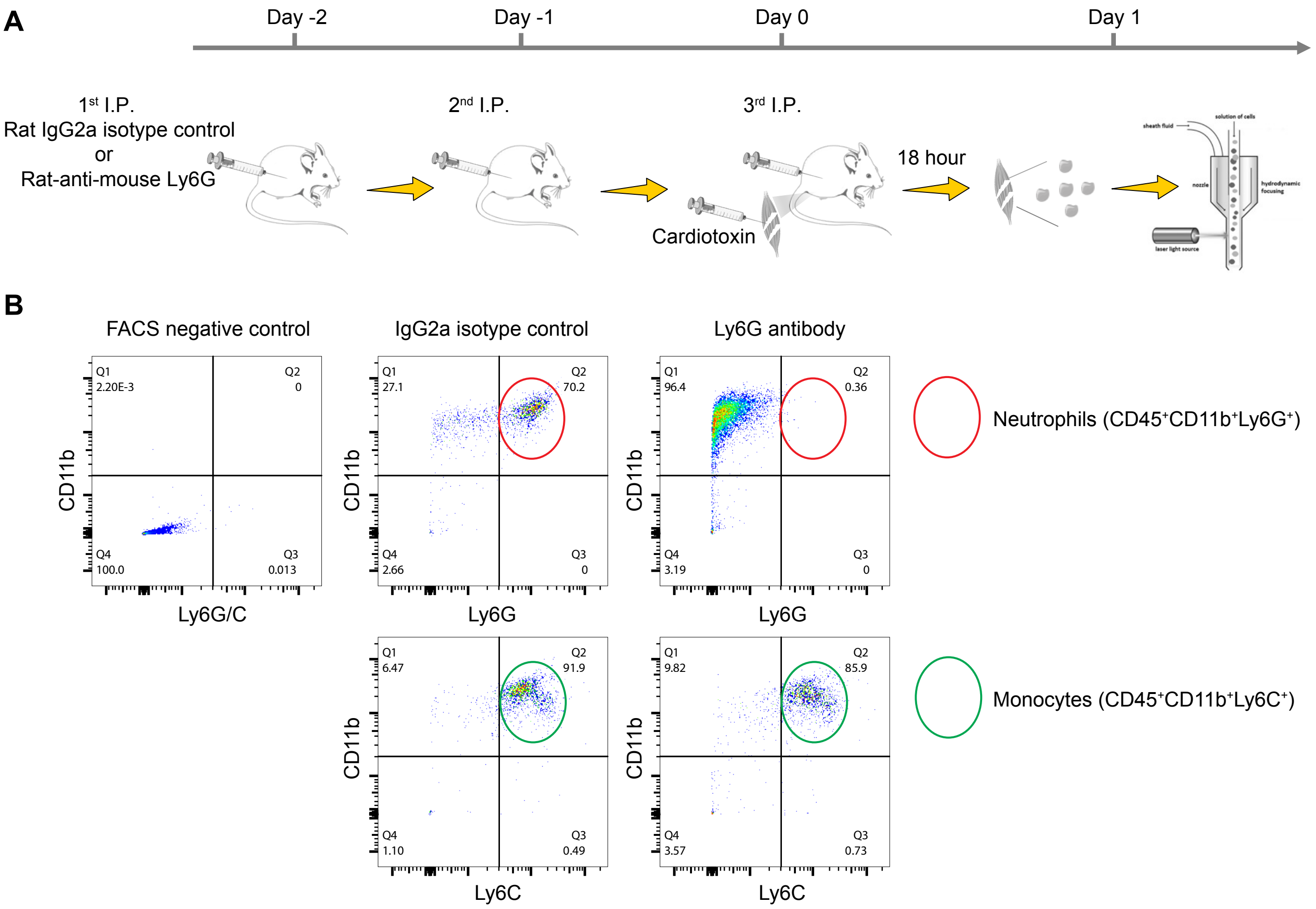
A
Day -2
Day -1
Day 0
Day 1
1st I.P.
Rat IgG2a isotype control
or
Rat-anti-mouse Ly6G
2nd I.P.
3rd I.P.
Cardiotoxin
18 hour
B
FACS negative control
IgG2a isotype control
Ly6G antibody
Q1 2.20E-3
Q2 0
Q4 100.0
Q3 0.013
CD11b
Ly6G/C
Q1 27.1
Q2 70.2
Q4 2.66
Q3 0
Ly6G
Q1 96.4
Q2 0.36
Q4 3.19
Q3 0
Neutrophils (CD45+CD11b+Ly6G+)
Q1 6.47
Q2 91.9
Q4 1.10
Q3 0.49
Ly6C
Q1 9.82
Q2 85.9
Q4 3.57
Q3 0.73
Monocytes (CD45+CD11b+Ly6C+)

### Figure S10

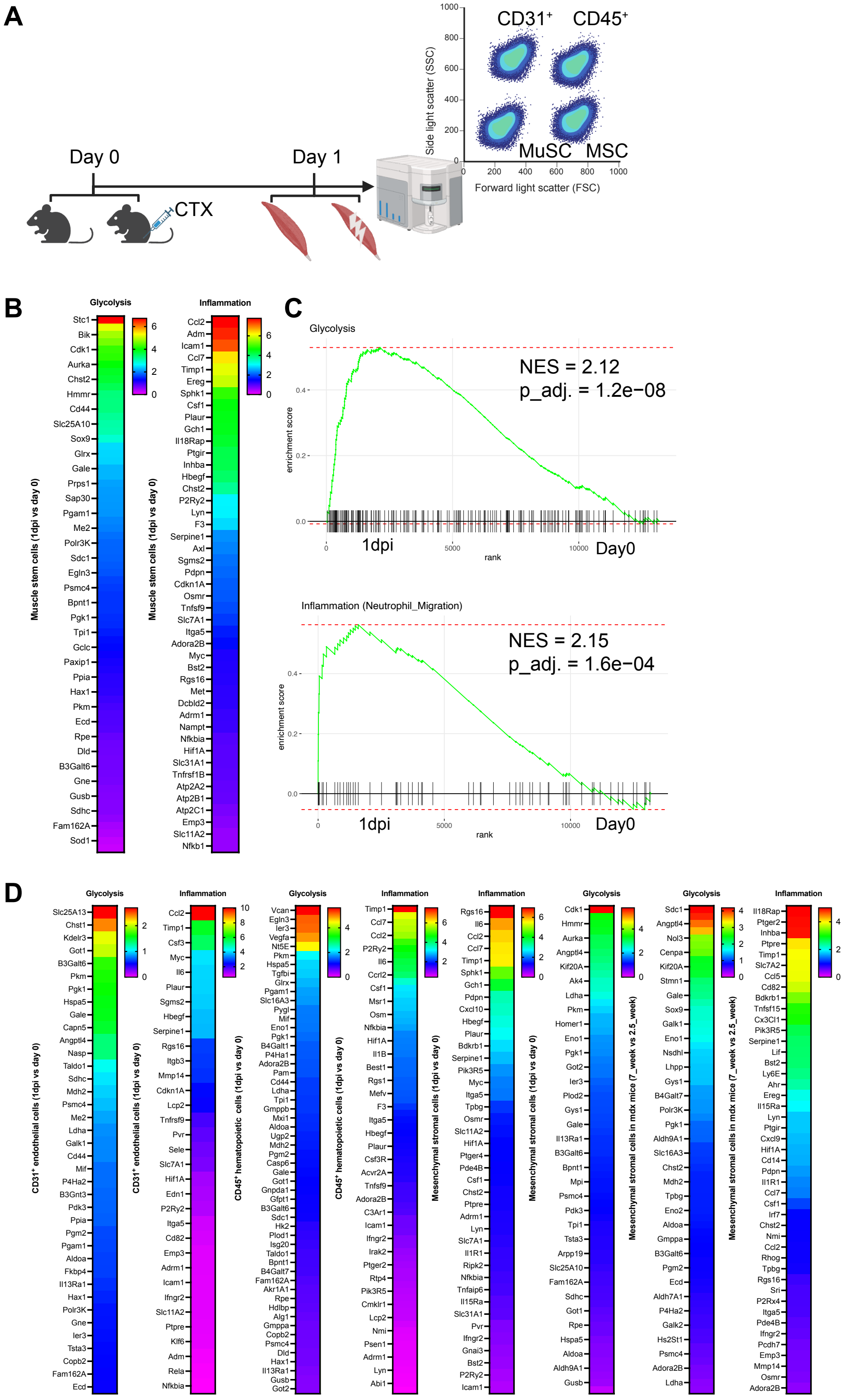
A
Day 0
Day 1
CTX
CD31+
CD45+
MuSC
MSC
Side light scatter (SSC)
Forward light scatter (FSC)
B
Glycolysis
Inflammation
Muscle stem cells (1dpi vs day 0)
C
Glycolysis
NES = 2.12
p_adj. = 1.2e−08
enrichment score
rank
1dpi
Day0
Inflammation (Neutrophil_Migration)
NES = 2.15
p_adj. = 1.6e−04
D
Glycolysis
Inflammation
CD31+ endothelial cells (1dpi vs day 0)
CD45+ hematopoietic cells (1dpi vs day 0)
Mesenchymal stromal cells (1dpi vs day 0)
Mesenchymal stromal cells in mdx mice (7_week vs 2.5_week)

#### SUPPLEMENTAL FIGURE LEGENDS

**Figure S1 The hypoxic niche of SCs in early stages of muscle regeneration**

(A) Western blot analysis showing the expression levels of HIF1α and HIF2α expression in hemostatic and cardiotoxin-injured TA muscles, from day 1 to day 5 post-injury. DPI, day post-injury.

(B) The time-course expression of canonical HIF1α target genes in SCs during cardiotoxin-induced muscle regeneration. The transcriptional data was sourced from a publicly available microarray dataset from a previous study[1].

(C) A diagram illustrating the bidirectional conversion of pyruvate to lactate mediated by lactate dehydrogenases.

(D) The temporal expression patterns of *Ldha* and *Ldhb* in SCs during cardiotoxin-induced muscle regeneration. The transcriptional data was retrieved from a publicly available microarray dataset of a previous study[1].

**Figure S2 Transcriptomic analysis of SCs during muscle regeneration**

(A) A volcano plot showing significantly differentially expressed genes (adjusted $p$-value < 0.05) in ASCs at 1 dpi compared to QSCs.

(B) The temporal expression profile of *Parp1* in SCs during cardiotoxin-induced muscle regeneration. Transcriptional data was retrieved from a publicly available microarray dataset of a previous study[1].

**Figure S3 Skeletal muscle single-cell RNA sequencing analysis**

(A) A dot plot illustrates the normalized expression levels of selected marker genes across various muscle cell types. The size of each dot indicates the proportion of cells within a specific cell type expressing the gene, while the color of the dot denotes the gene expression level, with blue indicating high expression and white indicating low expression.

(B) A volcano plot reveals genes that were significantly differentially expressed (adjusted $p$-value < 0.05, logFC > 0.25) between the type 1 and 2 neutrophil populations.

(C) Pathway enrichment analysis, based on gene ontology biological processes (GOBP) and Elsevier Pathway Collection, for genes significantly upregulated (adjusted p-value < 0.05, logFC > 0.25) in type 1 and type 2 neutrophils, respectively.

**Figure S4 Characterization of a mouse strain expressing cytosolic LbNOX**

(A) Schematic of the gene targeting strategy used to create the cytoLbNOX-expressing mouse strain (LbNOXfx/fx). This strategy involves the insertion of a loxP-flanked STOP codon upstream of the cytoLbNOX, followed by an internal ribosome entry site (IRES) concatenated enhanced green fluorescent protein (EGFP) sequence.

(B) Representative fluorescence images showing EGFP expression in primary myoblasts from the hindlimb muscles of LbNOXfx/fx mice, following induction with Cre recombinase-expressing adenovirus. Scale bar, 200 μm.

(C-E) Relative quantification of intracellular $NAD^+$ (C), NADH (D) levels, and the NADH/NAD+ ratio (E) in control versus cytoLbNOX-overexpressing myoblasts. * $p < 0.05$, *** $p < 0.001$ by Students' t test.

**Figure S5 Cytosolic $NAD^+$ regeneration is essential for the hypoxia-induced expression of immunomodulator genes**

(A) A schematic diagram illustrating the pharmacological blockade of $NAD^+$ regeneration in the conversion of pyruvate to lactate by inhibiting the catalytic activity of the enzyme LDHA.

(B) qPCR analysis demonstrating the expression levels of selected immunomodulator genes in wild-type myoblasts subjected to hypoxia (1% $O_2$) for 24 hours, both in the absence and presence of the lactate dehydrogenase inhibitor Gallofalvin. Myoblasts cultured under normoxia (21% $O_2$) served as the control group. * $p < 0.05$, *** $p < 0.001$ by One-Way ANOVA, comparing each group to myoblasts under normoxia conditions.

(C) qPCR analysis demonstrating the expression levels of *Ccl2* and *Itgb2* in *cytoLbNOX*-overexpressing and control myoblasts in the absence and presence of the lactate dehydrogenase inhibitor Galloflavin. * $p < 0.05$, *** $p < 0.001$ by One-Way ANOVA with the corrected post hoc multiple comparisons by Tukey.

(D) Western blot analysis revealing the levels of global PARylation, HIF1α, total and serine 536 phosphorylated NF-κB p65 subunit, with β-actin serving as the internal control, in wild-type myoblasts either cultured under normoxia or exposed to hypoxia for various durations.

**Figure S6 Circadian transcription activator Bmal1 regulates hypoxic induction of immunomodulator gene expression**

(A) Experimental design: Wild-type and *Bmal1*$^{-/-}$ myoblasts were subjected to 6 hours of hypoxia (1% $O_2$), with 3 replicates per condition. The RNA-sequencing data was derived from our previously published study[2].

(B) KEGG pathway enrichment analysis using genes significantly downregulated (adjusted *p*-value < 0.05) in *Bmal1*$^{-/-}$ compared to wild-type myoblasts under hypoxia conditions.

(C) A heatmap displays genes significantly (adjusted *p*-value < 0.05) downregulated and associated with the chemokine signaling pathway in *Bmal1*$^{-/-}$ versus wild-type myoblasts exposed to hypoxia for 6 hours. Color scale denotes log2 fold change.

**Figure S7 NicheNet analysis of interactions between MuSC and type 1 neutrophil populations**

(A) Heatmap showing NicheNet inferred receptors that express in the type 1 neutrophils and mediate the interaction with MuSC expressed ligands that are top-ranked in the ligand activity analysis.

(B) Heatmap displaying NicheNet inferred type 1 neutrophil-expressed active target genes of ligands that are top-ranked in the ligand activity analysis

**Figure S8 Analysis of gene expression in MuSC subclusters**

(A) A violin plot showing the expression profiles of established stage-specific marker genes across different MuSCs subclusters.

(B) A UMAP visualization of the transcriptomes of MuSC subclusters, categorized by days post-injury, providing a detailed view of the cellular heterogeneity and dynamics during muscle regeneration.

(C) Pseudotime analysis depicts single-cell expression trajectories of key myogenic regulatory factors, including *Pax7*, *Myod1*, and *Myog*, at ZT4 and ZT16, across the inferred myogenic lineages.

**Figure S9 Impact of Ly6G antibody injection on neutrophil infiltration at early muscle regeneration**

(A) Experimental design: wildtype C57BL/6 mice were subjected to transient neutrophil depletion through 3 consecutive intraperitoneal injections of a Ly6G antibody. For comparison, control mice received injections of a rat IgG2a antibody, which is of the same host species origin as the Ly6G antibody. After the last antibody treatment, all mice were subjected to intramuscular injections of cardiotoxin into the TA muscles. The presence of neutrophils and monocytes was evaluated via flow cytometry 18 hours post-injury.

(B) Representative FACS plots showing the gating strategy used to identify neutrophils ($CD45^+CD11b^+Ly6G^+$) and monocytes ($CD45^+CD11b^+Ly6C^+$) in the TA muscles injured by cardiotoxin in mice treated with Ly6G and IgG2a antibodies.

**Figure S10 Glycolysis and inflammation gene upregulation in muscle resident cells under hypoxia conditions**

(A) A schematic overview of the bulk RNA-sequencing experiment conducted in a previously published study[3]. This analysis involved several cell populations, including MuSCs, $CD31^+$ endothelial cells, CD45+ hematopoietic cells, and mesenchymal stromal cells (MSCs) from both homeostatic muscles and muscles injured 1dpi , which were then subjected to bulk RNA-sequencing.

(B) Heatmaps displaying genes significantly (adjusted *p*-value < 0.05) upregulated and associated with the glycolysis and inflammation signaling pathways in ASCs at 1 dpi compared to QSCs, based on the RNA-sequencing data from the study described in (A). Color scale denotes log2 fold change.

(C) Fgsea plots showing the concurrent activation of glycolysis and neutrophil chemotaxis and migration signaling pathways in ASCs at 1 dpi relative to QSCs.

(D) Heatmaps displaying genes significantly (adjusted *p*-value < 0.05) upregulated and related to glycolysis and inflammation signaling pathways in $CD31^+$ endothelial cells, CD45+ hematopoietic cells, and MSCs in injured muscles at 1 dpi, in comparison to their counterparts in homeostatic muscles, as well as in MSCs from the muscles of *mdx* mice at 7 weeks of age to those at 2.5 weeks of age. Color scale denotes log2 fold change.
